# Atomic structure of wheat ribosome reveals unique features of the plant ribosomes

**DOI:** 10.1101/2023.05.22.541707

**Authors:** Rishi Kumar Mishra, Prafful Sharma, Faisal Tarique Khaja, Adwaith B. Uday, Tanweer Hussain

## Abstract

Ribosomes from plants have unique plant-specific features that may aid in rapid gene expression and regulation in response to changing environmental conditions due to their sessile nature. Here, we present high-resolution cryo-electron microscopy structures of the 60S and 80S ribosomes from wheat, a monocot staple crop plant (*Triticum aestivum*). We compare wheat ribosome with closely related ribosomes from a dicot plant and other eukaryotes from yeast to humans. While plant ribosomes have unique plant-specific rRNA modification (Cm1847) in peptide exit tunnel, Zinc-finger motif in eL34 is absent and uL4 is extended making an exclusive interaction network. We note striking differences in eL15-Helix 11 (25S) interaction network, eL6-Expansion segment 7 assembly and certain rRNA chemical modifications between monocot and dicot ribosomes. Among eukaryotic ribosomes, we observe that rRNA modification (Gm75) in 5.8S rRNA is highly conserved and a base flipping (G1506) in peptide exit tunnel, and these features are likely involved in sensing nascent peptide. Finally, we discuss importance of universal conservation of three consecutive rRNA modifications in all ribosomes for their interaction with A-site aminoacyl-tRNA.

## Introduction

Cellular protein synthesis is a fundamental process in all life forms. Ribosomes are giant molecular machinery that carry out protein synthesis in cells. These macromolecular machines are asymmetrical assemblies of ribosomal RNA (rRNA) and ribosomal proteins (RP). All ribosomes contain two subunits: a small subunit and a large subunit. The small subunit helps decode codons in the mRNA, while the large subunit performs the peptidyl transferase activity to form a growing polypeptide chain. In prokaryotes, the small subunit (30S) is composed of 21 RPs and one rRNA (16S rRNA), while the large subunit (50S) contains 33 RPs and two ribosomal RNA (23S and 5S rRNA) (Melnikov *et al*, 2012). During the evolution from bacteria to eukaryotes, the ribosomes have increased in size and complexity. In eukaryotes, the large subunit (60S) contains three rRNA (25S/28S rRNA, 5.8S rRNA and 5S rRNA) with 47 RPs, while the small subunit (40S) is composed of one rRNA (18S rRNA) and 33 RPs (Yusupova & Yusupov, 2014). Although the core structure of the ribosome is very similar across eukaryotes, key differences exist in the ribosomes of different eukaryotic organisms.

Outside the shell of strongly conserved rRNA structure lies the additional blocks of rRNA in eukaryotic ribosomes known as Expansion segments (ES). Across eukaryotes, ribosomes differ in the length and sequence of rRNA in ES, which are the hotspot of diversity in the ribosome and play essential roles in stress response, mRNA binding, co-translational protein folding, and ribosome biogenesis (Hariharan et al, 2022, Fujii et al, 2018; Shankar et al, 2020; Parker et al, 2018). Ribosomes also differ in RP extensions and interaction networks (Timsit *et al*, 2021) as well as the distribution of chemical modifications on rRNAs and RPs. (Decatur & Fournier, 2002; Wu *et al*, 2021; Natchiar *et al*, 2018; Matzov *et al*, 2020; Streit & Schleiff, 2021; Sloan *et al*, 2017). Previous high-resolution structures of ribosomes from different eukaryotic species have been extremely helpful in providing insights into these features (Yusupova & Yusupov, 2017; Natchiar et al, 2017; Hopes et al, 2022; Matzov et al, 2020; Hiregange et al, 2022; Cottilli et al, 2022).

Plants being sessile in nature, undergo rapid gene expression and regulation in response to changing environmental conditions (Merchante *et al*, 2017). Hence the translational machinery in plants is different from other eukaryotes as it has plant-specific features and multiple isoforms of eukaryotic initiation factors (eIFs) (Gallie, 2016). The plant ribosome is also unique in possessing multiple functional paralogs of all ribosomal proteins (Martinez-Seidel *et al*, 2020), extension in ribosomal proteins (uL4, uL24, eL6, eL19, uS2, uS5 and eS10) compared to yeast (Lan *et al*, 2022) as well as high density of chemical modifications on plant rRNA (Streit & Schleiff, 2021). A low-resolution structure of plant 80S ribosome is available (Gogala et al, 2014), however, chemical modifications and details of interaction networks can only be visualised at the atomic resolution map of a plant ribosome.

Among crop plants, wheat is one of the most important staple crops as well as a widely used system for understanding plant biology. Wheat is highly susceptible to fungal infections, which lead to a huge loss in productivity (Figueroa *et al*, 2018). A structural understanding of plant protein synthesis and its comparison with structures of fungal protein synthesis machinery would open the possibilities of the development of potential antifungal drugs for plant diseases. Moreover, biochemical studies to understand protein synthesis in plants have been performed using wheat germ extract (Metz *et al*, 1999; Park *et al*, 2004; Harbers, 2014; Toribio *et al*, 2019). Therefore, we determined structures of the 60S and 80S ribosomes at atomic resolution from wheat, a monocotyledon crop plant species, using single-particle cryo-electron microscopy (cryo-EM). While we were working on the structure of wheat ribosomes, the structure of ribosomes from a dicotyledon plant, *Solanum lycopersicum* (tomato) (Cottilli *et al*, 2022), was reported recently.

Overall, the general architecture of the wheat ribosome is similar to the recently reported tomato ribosome structure and the plant-specific unique features are also observed in the wheat ribosome structure. However, comparing the wheat ribosome structure to the tomato ribosome structure reveals striking differences between closely related plant ribosomes from a monocot (wheat) and a dicot (tomato) plant. Moreover, we also observe and report additional plant-specific features in wheat ribosome structures. Finally, we also discuss a universally conserved modification in all kingdoms of life and its significance.

## Results & Discussion

### I. Overall architecture of wheat ribosome

Cryo-EM maps at atomic resolution were obtained for the 60S ribosomal subunit and whole 80S wheat ribosome from 2 different data sets (**Supp Figure 1A & 1B**). The maps were refined to a global resolution of 2.65Å (**Supp Figure 1C**) for the 60S subunit and 2.71Å for the 80S (**Supp Figure 1D**). Further, focussed refinements of large and small subunits of the 80S yielded cryo-EM maps of 60S and 40S at 2.69Å and 2.88Å, respectively (**Supp Figure 1E & 1F**). A focussed refinement of only the 40S body helped obtain a resolution of 2.84 Å (**Supp Figure 1G**). The local resolution of the core of the 60S is at 2.5Å or better in both maps. The availability of two atomic-resolution maps of the 60S from two independent data sets allowed us to confidently model the rRNA chemical modifications unambiguously and validate their definite presence in the wheat germ tissue (**Table 1 & Supp Figure 2A**). For the 40S subunit, the body is at higher resolution compared to the 40S head, and the extremities, like the 40S beak and the left/right feet, are at lower resolution.

**Figure 1.**
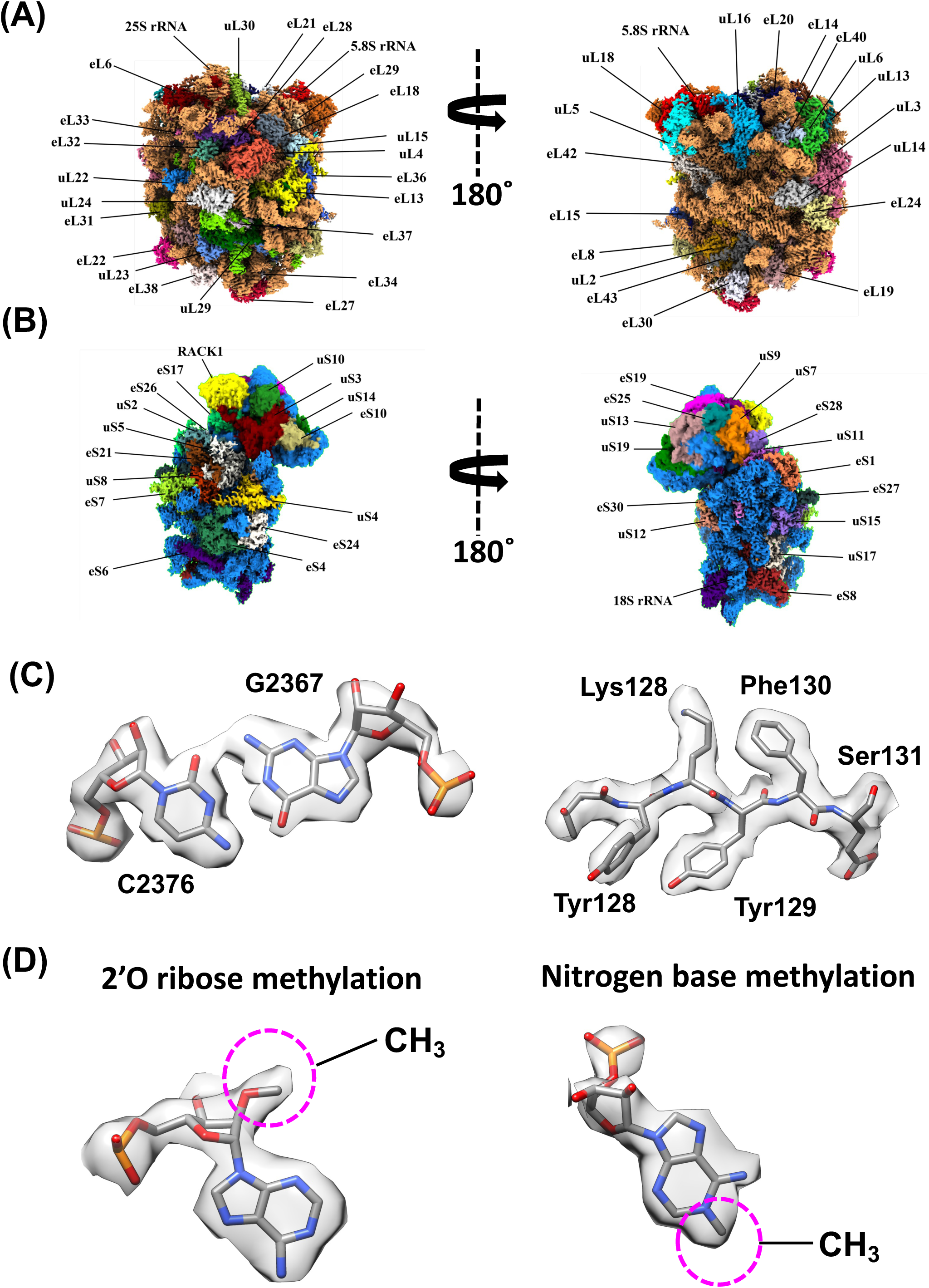
Overall structure of the wheat Ribosome. (A) Cryo-EM map of the 60S with 25S rRNA and protein coloured in the map with solvent face (left) & intersubunit face (right) (B) Cryo-EM map of the 40S with 18S rRNA and ribosomal proteins coloured distinctly in the map with the solvent face (left) & intersubunit face (right) (C) Model fit into the map for amino acid residues (left) and a nucleotide base pair (right), reflecting the quality of cryo-EM density in the structure. (D) Density display for 2’O ribose methylation (left) and Nitrogen base methylation (right)

**Figure 2:**
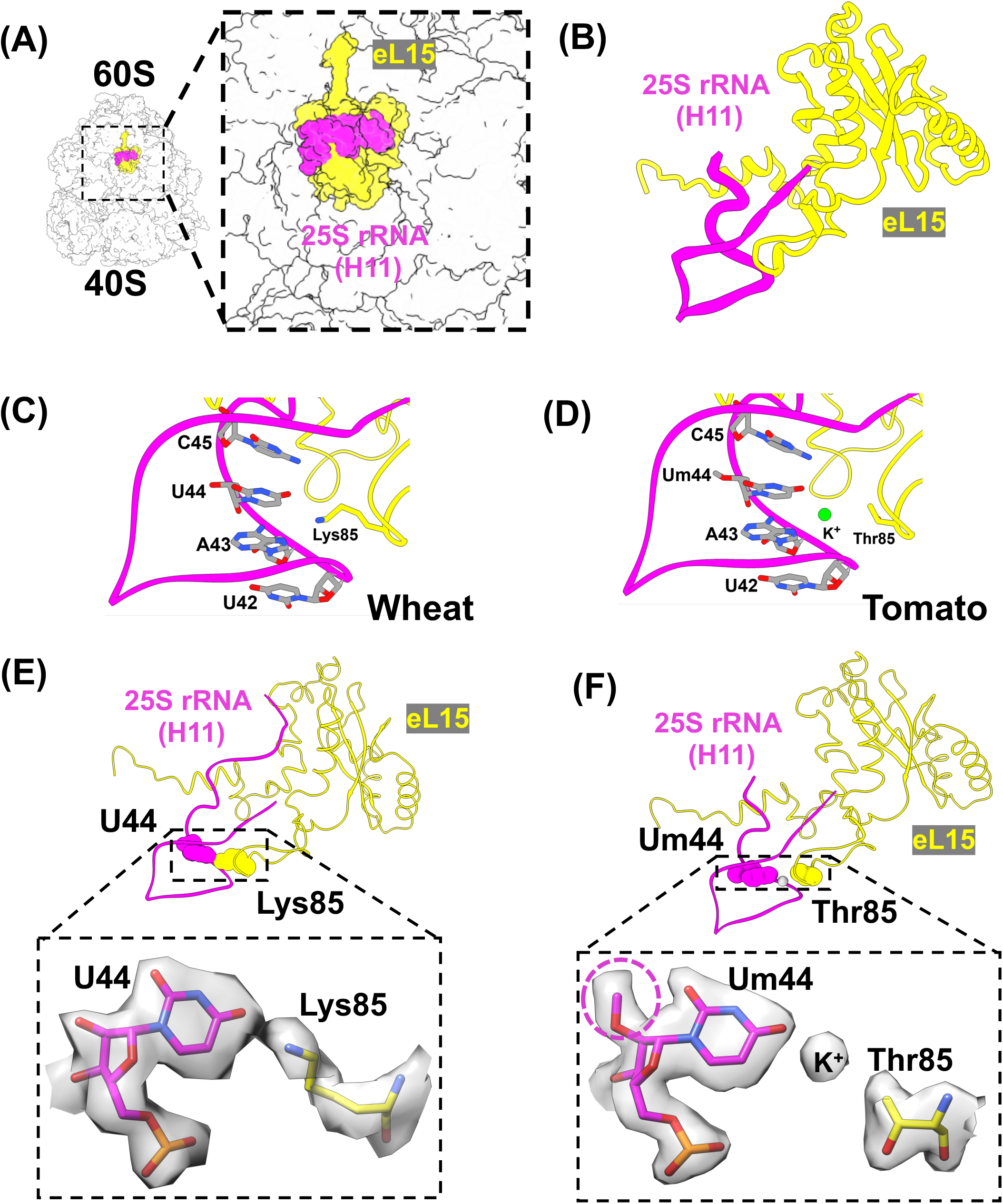
K85 of eL15 in monocots mimics the presence of K^+^ ion mediating interaction with H11 of rRNA. (A) eL15 is present in the large subunit and forms interaction with H11 of 25S rRNA (B) Interface of eL15 with H11 of 25S rRNA (C) In wheat (monocot) Lys85 of eL15 mediates interaction between eL15 and H11 of 25S rRNA (D) In tomato (dicot) the interaction is coordinated by the K^+^ ion that bridges Thr85 with U44 and the neighbouring bases in H11 (E) Density display representation of U44 in H11 of 25S rRNA showing the base to be unmodified in wheat (monocot) (F) The same base in the case of tomato (dicot) is 2’O-methylated, as represented in the density display diagram

We successfully built all ribosomal RNA (rRNA) (25S, 18S, 5.8S and 5S) and 79 RPs into the map (**Figure 1A & 1B**). The amino acid side chains and rRNA bases are modelled unambiguously (**Figure 1C & Table 1**). We searched for nucleotide modifications in rRNA based on previous biochemical studies on plant ribosomes (Wu *et al*, 2021; Azevedo-Favory *et al*, 2021; Streit *et al*, 2020; Sun *et al*, 2019; Streit & Schleiff, 2021a) and the recently reported tomato ribosome structure (Cottilli et al, 2022). We could directly visualize the density for 2’O ribose methylation and base methylation (**Figure 1D**). Pseudouridine, which has a similar geometry to uridine, was built into the map based on available reports for the modifications of plant rRNA (Sun *et al*, 2019; Streit & Schleiff, 2021; Cottilli *et al*, 2022). Thus, we modelled the chemical modifications in 25S, 18S, 5.8S and 5S rRNA (**Supp Figure 2A)**.

Overall, the general architecture of the wheat ribosome is similar to the recently reported tomato ribosome structure that is reflected by the similarity in rRNAs (rRNA) (25S, 18S, 5.8S and 5S) and RPs, post-translational modifications, and position of metal ions such as K^+^ ion and Mg^2+^. The plant-specific unique features reported in the recent structure of tomato ribosomes are also observed in the wheat ribosome structures obtained in this study **(Supp Figures 2B, 2C, 2D and 2E)**. Briefly, a direct contact between plant-specific 2’O methylated bases Am880, Cm2951 of 25S rRNA (wheat numbering is followed hereafter unless otherwise specified) and evolutionarily conserved methylated His246 of uL3 in the large subunit is also observed (**Supp Figure 2B**). Similarly, interactions of multiple methylated nucleotides Am821, Cm1844, Gm1846 (plant-specific) of 25S rRNA and Am43 (plant-specific) of 5.8S rRNA with the N-terminal region of eL37 (**Supp Fig 2C-2E**), which is involved in ribosome assembly, is also observed. Further, the overall chemical modification landscape of wheat ribosome is highly similar to that of the tomato ribosome.

In addition to the features mentioned above, we also observe additional plant-specific characteristics in wheat 60S and 80S structures obtained in this study that are not reported earlier (discussed below). Further, the availability of tomato (dicot) ribosome structure allowed us to compare the structures of two closely related ribosomes, i.e. from a monocot (wheat) and a dicot (tomato) plant, and report differences between the two.

### II. Differences between a monocot and a dicot plant ribosome

As mentioned above, the wheat and tomato ribosome structures are very similar to each other. However, upon extensive structural analysis, we observed striking differences between the two, which reflect differences in ribosomes from monocot and dicot plants. These differences are discussed in the subsections below.

#### a. Positively charged Lys sidechain in eL15 in monocots is replaced by Thr in dicots, and the charge is compensated by a K^+^ ion

eL15 is one of the RPs present in the large subunit (**Figure 2A**) and contributes to providing stability to the ribosome through its extensive interaction with the 25S rRNA. One of the interacting interfaces of eL15 is with H11 (helix 11) of 25S rRNA (**Figure 2B**). A striking difference in monocot ribosomes from that of the dicot ribosomes is a change in the last residue of the highly conserved VYGKPK motif in eL15. Monocots (including wheat) have conserved VYGKPK motif in eL15 with Lys at the 85^th^ position (**Figure 2C and Supp Figure 3A**), while in dicots (including tomato), the last residue is mutated to Thr (**Figure 2D and Supp Figure 3A)**. In wheat, the positively charged sidechain of Lys85 forms interactions with A43 and U44 in 25S rRNA and bridges eL15 with H11 of 25S rRNA (**Figure 2C**). Interestingly, in tomato, the residue Thr85 coordinates a K^+^ ion, which bridges the eL15 to rRNA bases in this region, as represented in **(Figure 2D).** This feature of Lys85 forming the bridge in monocots is similar to protozoa (**Supp Figure 3B & 3C**), while the presence of a cation (K^+^) in dicots is like fungi and metazoan where a cation mediates the interaction between H11 and eL15 (**Supp Figure 3D-3G**). Finally, a positively charged moiety is always present at this position irrespective of different amino acid residues in different organisms, which indicates that the interaction bridging H11 and eL15 might be crucial for the ribosome function. Further, we observe that Lys85 (in monocots) interacts with unmodified U44 of 25S rRNA (**Figure 2E**). Interestingly, dicots possess a plant-specific 2’O methylation on U44 (**Figure 2F**) which is absent in other ribosomes. It will be interesting to explore the role of this modification in dicots.

**Figure 3.**
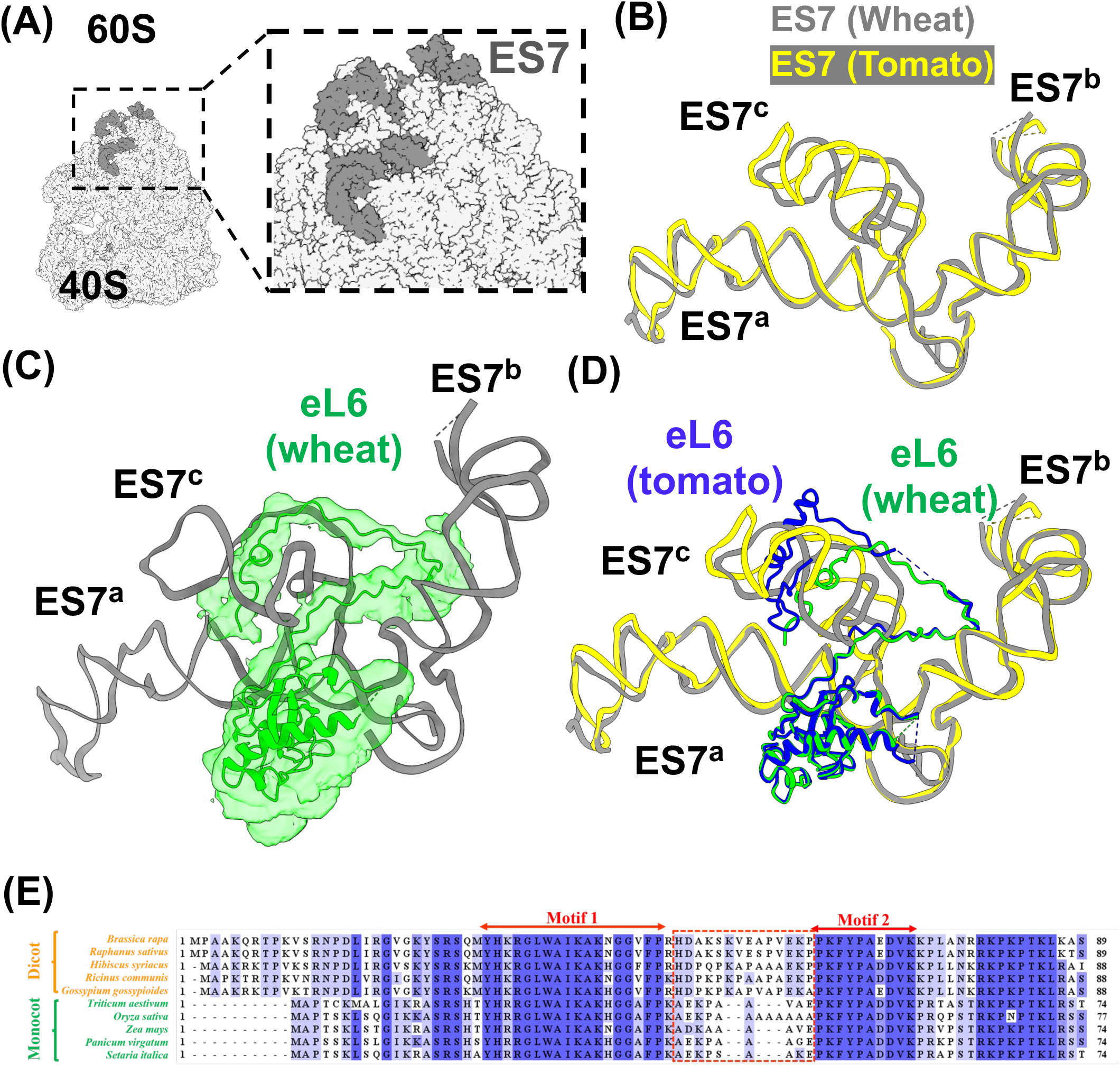
Difference in the conformation of ES7 between monocots and dicots. (A) ES7 depicted on the solvent interface of a large subunit of the wheat ribosome (B) Superposition of ES7 from wheat (grey) and ES7 from tomato (yellow) exhibits a conformation difference in ES7^c^ (C) Ribosomal protein eL6 intercalates and passes through the two strands of ES7^c^ in wheat as represented by the Cryo-EM density display of eL6 (D) Superposition of wheat (monocot) and tomato (dicot) ES7-eL6 structure shows the position of intercalation of eL6 N-terminal differs between wheat and tomato ribosome (E) Sequence alignment of the N-terminal portion of eL6 comparing monocot and dicot species where the conserved motifs are highlighted by a magenta line on the top and the linker between conserved motifs shown inside the dashed box

#### b. Difference in the conformation of ES7-eL6 assembly

The Expansion segment 7 (ES7), present on the solvent face of the large subunit **(Figure 3A),** is one of the most diverse expansion segments (M *et al*, 2022) in eukaryotes, which closely interacts with uL4, eL28 and eL6. ES7 binds to diverse factors involved in stress response, translational fidelity, and ribosome biogenesis (Gómez Ramos et al., 2016; Shedlovskiy et al., 2017). The structural superposition of ES7 from wheat and tomato show a difference in the conformation of ES7^c^ (**Figure 3B**). This branch of ES7 interacts with eL6, which is involved in evoking stress response in crop plants (Moin *et al*, 2020; Sahi *et al*, 2006; Pei *et al*, 2019; Islam *et al*, 2020). When we analysed the interaction between eL6 and ES7, we observed that the N-terminal tail (NTT) of eL6 in wheat ribosomes intercalates and passes through the two strands of the c loop of ES7 (**Figure 3C**), which is a feature present in plants and metazoan (**Supp Figure 4A & 4B**) and absent in case of fungi and protozoan (**Supp Figure 4C & 4D**).

**Figure 4.**
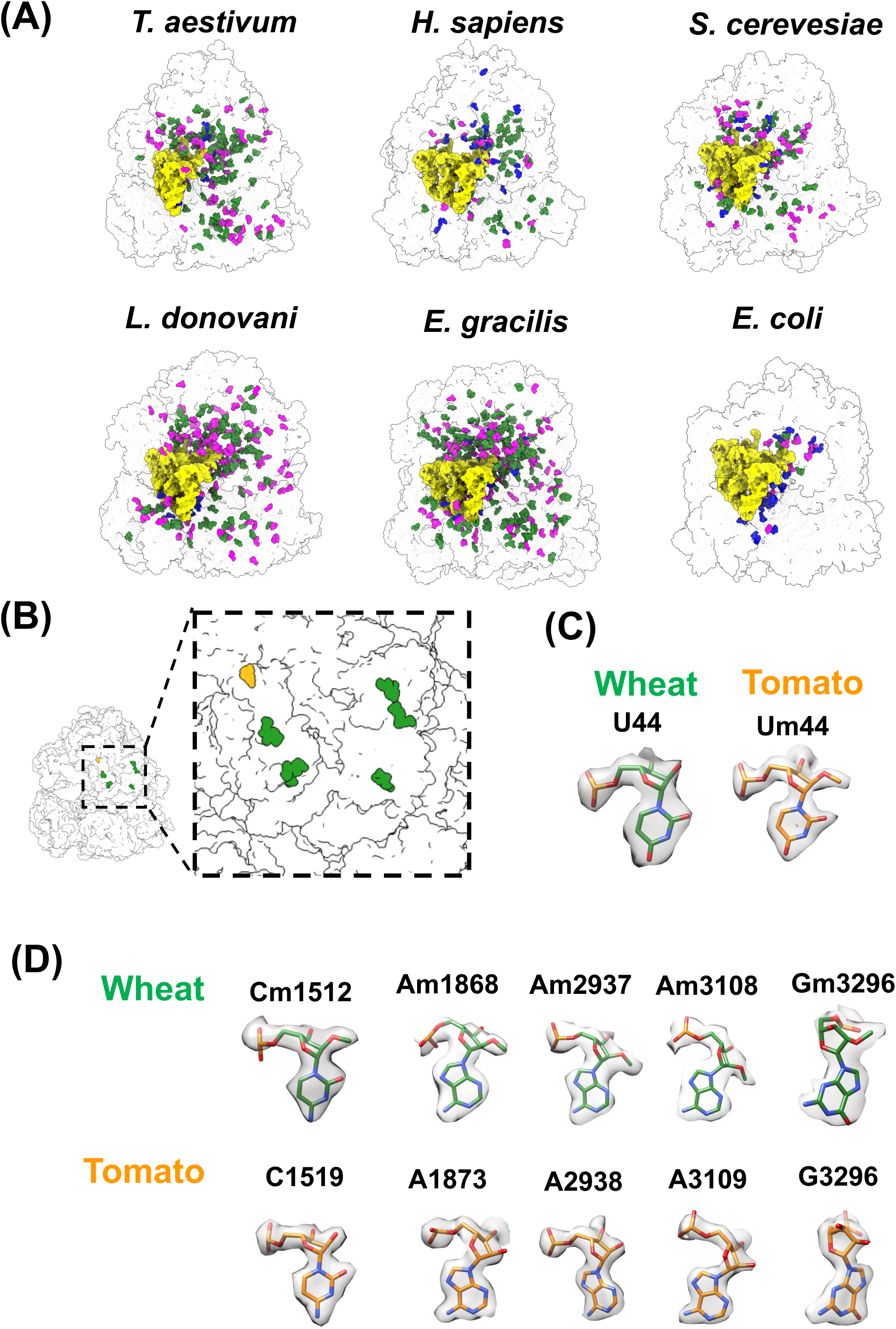
Difference in chemical modification of rRNA between wheat and tomato ribosomes. (A) Chemical modifications observed in ribosomes of different species, namely *T. aestivum* (this study), *H. sapiens* (PDB-ID: 6QZP), *S. cerevesiae* (4V88, modifications highlighted based on (Sloan *et al*, 2017)), *E. gracilis* (PDB-ID:6ZJ3), *L. donovani* (PDB-ID: 6AZ3), and *E. coli* (PDB-ID: 7K00). Plants show a higher density of chemical modifications compared to bacteria, yeast, and humans. (B) The position of 2’O ribose methylated residues on plant ribosomes, which are unique to wheat (green) or tomato (orange) (C) 2’O-methylation on U44 was observed only in tomato, and density for methylation is absent in the wheat ribosome map (D) A variety of 2’O-methylation, Cm1512, Am1868, Am2937, Am3108 and Gm3296, was observed at 2’ribose in wheat (top panel) while the density for the same was absent in the tomato ribosome (bottom panel)

Interestingly, the position of eL6 intercalation differs in wheat and tomato (**Figure 3D**), and this is likely because of the presence of a shorter linker between two conserved motifs in eL6 in wheat (Figure 3E). The first conserved motif RGLWAIKAKN/HGG (**Motif 1**), interacts with ES7^c^, while the second conserved motif PKFYPAD/EDVK (**Motif 2**), interacts with ES7^b^ and nearby RPs. The linker between the two motifs is shorter in monocots in comparison to the dicots (**Figure 3E**), which leads to the observed difference in the position of intercalation of NTT of eL6 through ES7^c^ between wheat and tomato ribosome.

Residues of eL6 interacting with ES7^c^ are conserved across all the plant species (**Supp Figure 4E**). To visualise the interaction between conserved residues of eL6 with ES7^c^, we used the structure of tomato ribosome as the resolution in this region of the wheat ribosome is not high enough to observe atomic interactions. Within the conserved stretch in Motif1, we observe that the Trp31 and a pair of consecutive glycine (Gly38Gly39) are present only in plants. Trp31 forms plant-specific stacking interaction with G595 of ES7^c^ (**Supp Figure 4F**), while Gly38Gly39 dipeptide appears to facilitate the bending of eL6-NTT for passing through ES7^c^ (**Supp Figure 4G**). Despite the difference in the conformation of ES7^c^-eL6 assembly in monocots and dicots, the residues involved in the intercalation of eL6 (NTT) through ES7^c^ are conserved in plants which might be crucial for maintaining ES7 in a conformation conducive to binding the factors involved in stress response, translation fidelity and ribosome biogenesis.

#### c. Differences in rRNA modifications

Comparing the landscape of chemical modifications of rRNA in wheat with the ribosomes of other organisms **(Figure 4A)** shows that plants have a higher density of rRNA modifications compared to yeast and human ribosomes but lesser compared to protozoa that harbour fragmented ribosomal RNAs. The high number of chemical modifications in protozoa was hypothesized to stabilize the fragmented rRNA structure, likewise, the high number of rRNA modifications in plants might help stabilize the rRNA during varying environmental conditions.

Among plants, we observe differences in the occurrence of chemical modification of rRNA, even in the closely related monocots and dicots (**Figure 4B**). As discussed earlier in tomato (dicot), U44 of 25S rRNA is 2’O-methylated, while the same modification is absent in the wheat (monocot) ribosome structure (**Figure 2E, 2F and 4C**). Interestingly, this modification is also absent in yeast as well as mammals. With available information from structures ((Cottilli *et al*, 2022) and this work) and mass-spectrometry as well as RiboMethSeq reports (Cottilli *et al*, 2022; Azevedo-Favory *et al*, 2021), we are unable to find any other rRNA modifications specific to dicots only. Similarly, the wheat ribosome shows the presence of 2’-O ribose methyl groups on a few nucleotides of the 25S rRNA (Cm1512, Am1868, Am2937, Am3108 and Gm3296), while these modifications are absent in the tomato ribosomes **(Figure 4D)**. None of these modifications is seen in fungi, while in the case of mammals, Cm1512 and Am1868 are visualized, and Um44, Am2937, Am3108 and Gm3296 are absent. As these modifications are present towards the peripheral region of the large subunit, they may be transient in nature or tissue-specific, but the exact role of these modifications needs further exploration.

### III. Other Plant-specific features in ribosomes

Apart from the plant-specific features reported in tomato ribosome (Cottilli *et al*, 2022) structure, we also observe additional plant-specific features, which we describe in subsequent subsections below. These features are observed in wheat ribosomes in our study and recently reported ribosome structure from tomato. However, these features were not discussed in the recent paper by Cottilli *et al*.,2022. We will mention the features of wheat as a reference for plant ribosomes. As mentioned earlier, we observe these features in both the 60S as well as 80S maps in our study.

#### a. Plants-specific 2’O methylation in Peptide Exit Tunnel (PET)

Besides the chemical modifications unique to monocots and dicots, we visualize the plant-specific modifications that are present in both. We visualize a plant-specific 2’O-ribose methylation of a conserved Cytidine (Cm1847) present in the peptide exit tunnel (PET) (**Figure 5A**). Cm1847 is present very close to the modelled nascent peptide in PET and can directly interact with it (**Figure 5B**). Further, Cm1847 interacts with the loop of uL22 that forms constriction in the PET of all the eukaryotes (**Figure 5C**). Unmethylated C1847 can form hydrogen bonds with uL22 residues R129 and Y131, while the 2’O methylated Cm1847 in plants do not form these hydrogen bonds (**Figure 5D**). As these amino acid residues are universally conserved (**Figure 5E**), the role of 2’O-methylation of conserved C1847 in plants is not clear and needs further exploration. As Cm1847 do not lead to any structural change in and around PET, we hypothesize that the 2’O-methylation of C1847 in plants might protect it from endonuclease during the early stage of ribosome biogenesis and thus might have a role in providing stability to rRNA in plants.

**Figure 5.**
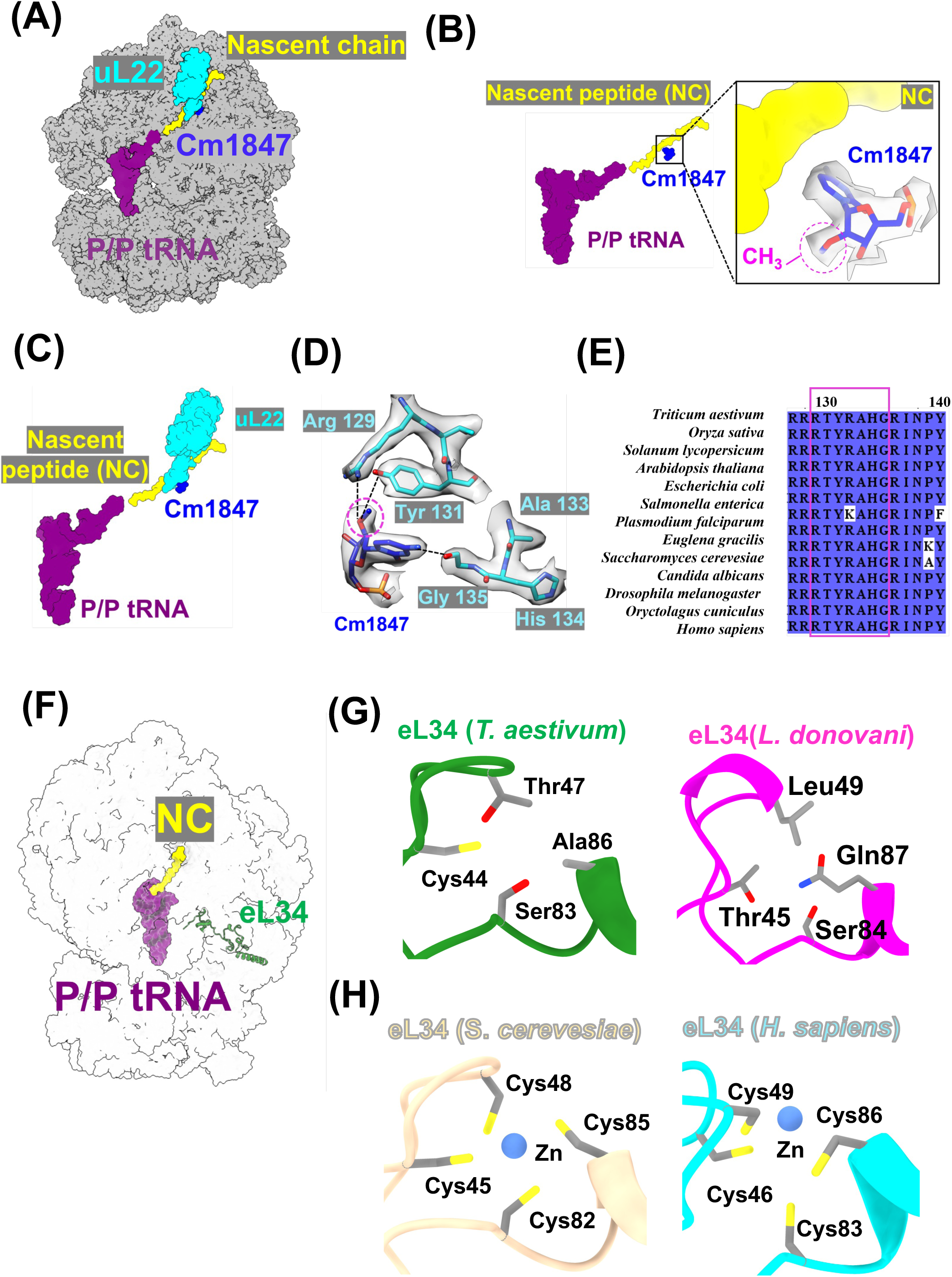
Plant-specific chemical modification present in the peptide exit tunnel and enigmatic absence of Zn ion in eL34 of plants (and protozoa) (A) Position of plant-specific 2’O ribose methylation, Cm1847, with respect to PET and uL22 on 60S (B) Interaction of Cm1847 with modelled nascent peptide chain where PDB ID – 7QWR has been used for modelling the nascent chain in the wheat ribosome (C) The modification Cm1847 forms direct contact with the portion of uL22 loop that forms constriction in the peptide exit tunnel (D) Methyl group on Cm1847 makes Van der Waals interaction with the uL22 residue R129 and Y131 (E) Sequence comparison of uL22 ribosomal protein where the residues forming constriction in peptide exit tunnel is highlighted in the magenta box (F) Ribosome depicting the position of eL34 (in green) on the large subunit (G) A close-up view of the Zinc-finger motif in eL34 of wheat representing the absence of Zn metal in eL34 of plants (*T. aestivum*) and protozoa (*L. donovani*) (H) Zoomed view of Zinc-finger motif in eL34 of representative organisms of other species (fungi (*S. cerevesiae*), and mammals (*H. sapiens*)

#### b. Absence of conserved Zinc-finger motif of eL34 in plant and protozoa

Zinc-finger motifs bind to DNA and are present in many transcription factors in all domains of life. Zinc-finger motifs are also present in ribosomes of all species, from prokaryotes to eukaryotes. However, not much is known about the role of these motifs in the ribosome. eL34 is one of the Zinc-finger containing RP, which is present in the large subunit of the ribosome (**Figure 5F**). Our structure shows an absence of density for Zinc in the Zinc-finger motif of the eL34 (**Figure 5G**), which agrees with the recently reported structure of tomato ribosome (Cottilli *et al*, 2022). We performed sequence analysis of eukaryotes from protists to mammals and observed the absence of the conserved cysteines of the Zinc-finger motif in eL34 in plants (**Supp Figure 5A**). These conserved cysteines of the Zinc-finger motif in eL34 are also absent in photosynthetic protists, e.g., Euglena and other species of protozoa in class sarcomastigophora (**Supp Figure 5A**). Further, we investigated the available high-resolution structure of ribosomes from different species of protozoa and observed a similar absence of Zn ion in eL34 (**Figure 5G**). Thus, the absence of Zn ion in eL34 is not specific to plants as proposed earlier (Cottilli *et al*, 2022). On the other hand, the Zn ion is bound to eL34 in yeast and mammals (**Figure 5H**). Superposition of the eL34 structure with and without the Zinc-finger motif exhibits no difference in the overall conformation of the protein (**Supp Figure 5B**), ruling out the role of the Zn ion in the conformation of eL34. As these motifs are well conserved in yeast and higher eukaryotes, we wondered why these are lost in protozoa and plants.

Early work on ribosomal Zinc-finger motifs has shown that the first cysteine in the motif is crucial for translation, while the following three cysteines and Zinc are dispensable (Rivlin *et al*, 2000). Additionally, the presence of a positive stretch of amino acid sequence with conserved aromatic residues in between the cysteines forming the Zinc-finger motif is sufficient for the protein synthesis (Rivlin *et al*, 2000; Dresios *et al*, 2002). In line with this, plant eL34 has conserved cysteine at the first position as well as a basic amino acid-rich residue intervening sequence (**Supp Figure 5A)** where the positively charged amino acid forms multiple contacts with rRNA as listed in (**Table 5)**. These interactions are sufficient for stabilizing eL34 on the ribosome, making the zinc-finger motif dispensable for its ribosome-specific function. Besides this, the extra-ribosomal activity of different RPs has also been discovered (Warner & McIntosh, 2009), which is more prevalent in the case of plants (Xiong *et al*, 2021). Therefore, we hypothesize that the plants and protozoans once possessed the Zinc-finger motif in eL34, which might have been essential for its yet-to-be-understood extra-ribosomal activity. During evolution, the extra-ribosomal action was taken over by some unknown factor or was no longer needed in these species; therefore, the Zinc and, thus, the dispensable cysteines were lost from plant eL34.

#### c. uL4 C-terminal tail shows plant-specific interactions

The eukaryote-specific C-terminalα-helical extension of uL4 makes numerous contacts with the surface RPs which includes uL30, eL20, eL18 and rRNA like ES7, and mutations in the C-terminal extension of uL4, show defects in growth as well as 60S subunit formation in yeast (Stelter *et al*, 2015). Plants have longer C-terminal tails (CTT) of uL4 compared to yeast (**Supp Figure 6A**). In the wheat ribosome, the extended C-terminal residues of uL4 make extensive interaction with eL20, eL21 and h43 and thus stabilize them (**Figure 6A**). The uL4 residues involved in these stabilizing interactions (**Figure 6B-6E)** are highly conserved in plants (**Supp Figure 6B**), suggesting that the interactions are plant specific. Stabilization of these interactions is important as the deletion of uL4 CTT leads to a significant growth defect at higher temperatures (37 °C) in yeast (Stelter *et al*, 2015). Therefore, it is likely that the longer uL4 CTT in plants interacts with the surrounding RPs and rRNA to further stabilize the local network of interactions, which might help plants survive through varying stress and environmental conditions. Notably, in humans, these RPs are stabilized by an expansion segment ES7L instead of uL4 CTT (**Figure 6F**). The uL4 CTT in humans, which is longer than the plants, appear flexible as it is not captured in any structural study to date (Natchiar et al, 2017; Hopes et al, 2022). This suggests that, like lower eukaryotes, plants have evolved a different mechanism of uL4 stabilization compared to humans.

**Figure 6.**
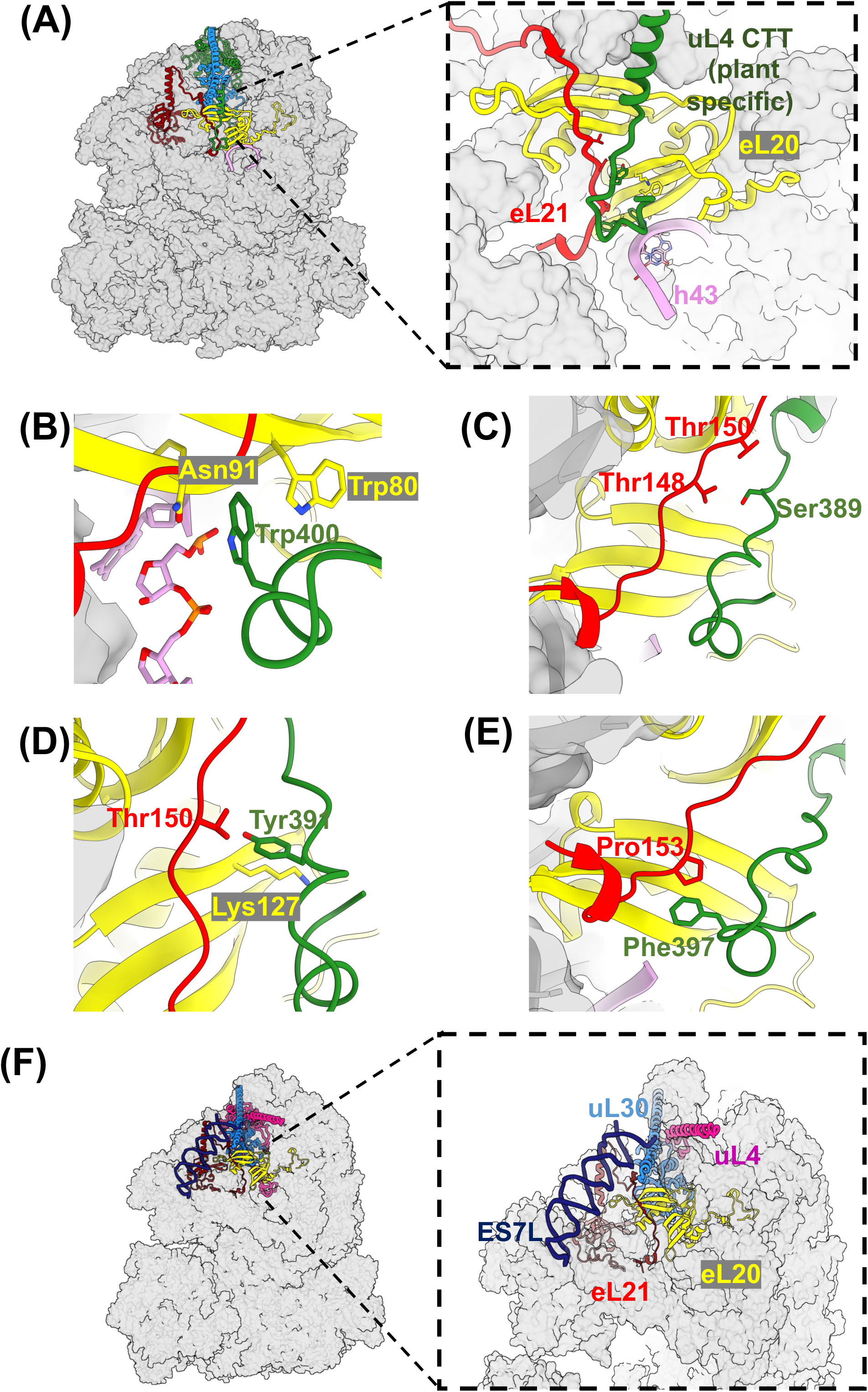
Ribosomal protein uL4 in wheat forms plant-specific conserved interactions at its C-terminal tail. (A) Location of uL4 (green), eL20 (yellow), eL21 (red) and uL30 (blue) on wheat 80S ribosome and the zoomed view of uL4 CTT on ribosome showing its contact with eL21, eL20 and h43 (B) Close-up view of the region surrounding CTT of uL4 highlighting plant-specific interactions in uL4 CTT, e.g., Trp400 of wheat uL4 interacts with Trp80 and Asn91 of eL20 and with the phosphate backbone of h43 (C) Ser389 of wheat uL4 interacts with Thr148 and Thr150 of eL21 (D) Tyr391 of wheat uL4 interacts with Thr150 of eL21 and Lys127 of eL20 (E) Phe397 of wheat uL4 interacts with Pro153 of eL20 (F) Human ES7L (dark blue) superimposed on wheat 80S. Adjacent to it is the zoomed view showing the stabilization of human uL4 (pink), eL20 (yellow), eL21 (red) and uL30 (blue) by ES7L

### IV. Conserved features in eukaryotic ribosomes

Besides the findings unique to plant ribosomes, we noticed other interesting features in the active centres of wheat ribosome structure. We observe that G1506 lining the PET is flipped out into the exit tunnel (**Figure 7A**). This flipped conformation was also observed in thermophilic fungi, *Chaetomium thermophilum* where the flipping of corresponding nucleotide has been proposed to create a third constriction in PET for sensing the nascent peptide in the tunnel (Kišonaitė *et al*, 2022). However, in wheat 60S as well as 80S maps, G1506 is flipped out even in the absence of a nascent peptide in the exit tunnel. Notably, a similar flipping conformation is also present in tomato ribosomes where again the nascent peptide is absent (Cottilli *et al*, 2022). This suggests that the nascent peptide is not required for the flipped conformation of G1506, however the flipping might facilitate stabilization of the peptide in the PET of plants and thermophilic fungi. Thus, both plants and thermophilic fungi share a similarity in the mechanism of peptide stabilization in the exit tunnel.

**Figure 7.**
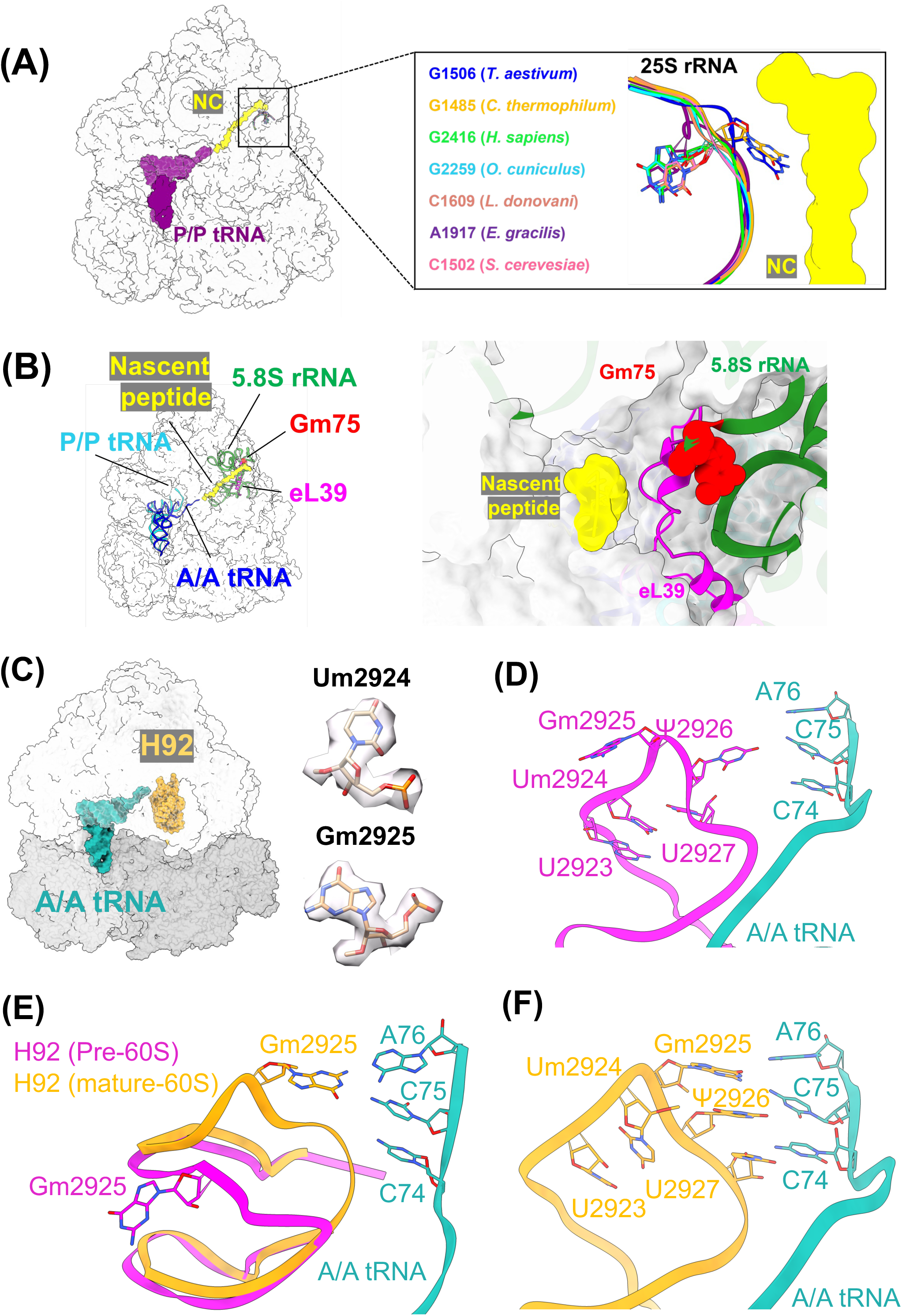
Features of plant ribosome similar to other eukaryotes and potential role of universally conserved triplet Um2924Gm2925Ψ2926 in helix 92 of 25S rRNA in tRNA accommodation. (A) A bulged-out conformation of G1506 in plants and its comparison with other species listed in the figure (PDBs used 6QZP (*H. sapiens*), 7OLC (*C. thermophilum*), 7O7Y (*O. cuniculus*), 6AZ3 (*E. gracilis*), 6ZJ3 (*L. donovani*), 4V88 (*S. cerevesiae*) and this work) (B) Location of the highly conserved Gm75 nucleotide near the exit tunnel and its direct binding with nascent peptide sensing protein eL39, where the nascent peptide is modelled using 7QWR (C) Position of H92 with respect to the A-site tRNA on the ribosome (D) A distorted conformation of the nucleotide in H92 near the triplet in pre-60S (PDB ID: 7UG6) showing a modelled A-site tRNA (modelled using PDB 6ZJ3) is unsuitable for forming interactions with the acceptor arm of tRNA, wheat numbering for rRNA bases have been used for clarity (E) A conformational transition of H92 during 60S subunit maturation brings the tip of H92 closer to the position of the acceptor arm of tRNA (F) The 2’O-methylated Um2924 and Gm2925 adopt a planar geometry of the nucleotides facilitating multiple stacking interactions with the neighbouring bases while Ψ2926 makes additional hydrogen bonds with the nucleotides

The 5.8S rRNA is also modified in all eukaryotic species. We note that despite the difference in the pattern of distribution of modification across species (**Supp Figure 7A**), 2’O-ribose methylation of G75 is highly conserved (**Supp Figure 7B & 7C**) (except in yeast). We observe the density for methylation of G75 of 5.8S rRNA in both 60S as well as 80S maps, which agrees with the recent structure from tomato (Cottilli et al, 2022). We notice that Gm75 directly contacts eL39 (**Figure 7B**), an RP involved in sensing the nascent peptide in the exit tunnel, which might be the reason for the conservation of this 5.8S rRNA methylation in eukaryotes.

### V. Universally conserved stretch of rRNA modifications in Helix 92 (H92) of 25S rRNA

We observe three consecutive modified nucleotide Um2924Gm2925Ψ2926 in H92 of the wheat ribosome in both 60S as well as 80S maps (Figure 7C). This triplet interacts with the CCA end of the aminoacyl-tRNA at the A-site (Kim & Green, 1999). Analysis of high-resolution structures and other biochemical studies on rRNA modifications of different species (Matzov et al, 2020; Natchiar et al, 2018; Cottilli et al, 2022) showed that the triplet of modified nucleotide Um2924Gm2925Ψ2926 is conserved throughout from bacteria to mammals including the organisms with highly atypical ribosome with fragmented rRNA, e.g., *Euglena gracilis* (**Supp Figure 8A**).

A recent report shows the conserved methylated Guanine Gm2922 (Yeast residue corresponding to wheat Gm2925) in helix92 to be essential for ribosome biogenesis in Yeast (Yelland *et al*, 2022). Therefore, we compared the helix92 in pre-60S (PDB ID 3JCT) with our structure of mature 60S and observed a distorted conformation of the modified nucleotide triplet in helix 92 in the case of pre-60S owing to interaction with ribosome biogenesis factors (**Figure 7C & 7D**) that might be crucial for biogenesis (Wu *et al*, 2016). This distorted conformation is not suitable for interaction with the CCA arm of aminoacyl-tRNA at the A-site in the mature ribosome (**Figure 7D**). During the final step of ribosome maturation, helix 92 undergoes a conformational change where the tip of H92 moves towards the A-site (**Figure 7E**). Recent reports show that the 2’O ribose methylation with C3’endo sugar pucker stabilizes the base’s planar conformation, which facilitates the additional stacking interactions to stabilize the rRNA (Natchiar *et al*, 2018). Analysis of our structure of mature wheat ribosome shows a similar 3’ endo conformation of the modified nucleotides Um2924Gm2925Ψ2926 in Helix 92 (**Supp Figure 8B**).

Therefore, we reason that methylation in Um2924 and Gm2925 facilitates the transition of distorted conformation of the nucleotides in pre-60S into a planar conformation in the mature 60S (**Figure 7F**). The additional hydrogen bonding by Ψ2926 as well as multiple stacking interactions, stabilizes the triplet in a planar stacked conformation suitable to base-pair with tRNA. This hypothesis is further strengthened by the structure of the yeast ribosome devoid of pseudouridylation (Zhao et al, 2022), where we observe a non-planar conformation owing to the lack of stacking with the neighboring nucleotides (**Supp Figure 8C**). This disturbed conformation has diminished interaction with tRNA. Thus, the planar stacked conformation of the Um2924Gm2925Ψ2926, stabilized by the modifications, facilitates its interaction with the CCA arm of the A-site tRNA in the mature ribosome (**Supp Figure 8D**), helping in the proper accommodation of aminoacyl-tRNA in the A-site and hence explaining the remarkable conservation of these modifications across evolution.

## Concluding remarks

In summary, we determined the atomic resolution structures of the 60S and 80S ribosomes purified from wheat germ extract. Wheat is a crop plant that belongs to the grass family (along with barley, rice, and maize) and is one of the largest consumed staple foods across the globe. It is highly prone to infection by pathogens like rust fungi, bacteria, plant viruses, and insects leading to huge economic losses (Singla & Krattinger, 2016). Understanding the unique features of the protein synthesis machinery of such important crop plants can help devise strategies to fight against these pathogens. Here we highlight differences between wheat ribosomes with other eukaryotic organisms like humans, yeast, and closely related tomato ribosomes. This study also provides a reference model for structural studies in plant translation as well as for carrying out structure-based evolutionary studies. Overall, we visualize the detailed structure of wheat ribosome as well as the features specific to plants which enhances our understanding of important similarities as well as the diversity of these macromolecular machines across species in the evolution.

## Material and methods

### Purification of wheat 80S ribosomes

For the purification of the ribosome, prechilled 100g of commercial wheat germ was blended into fine powder in a mixer grinder by giving five bursts for 15 seconds in liquid Nitrogen and resuspended in 200 mL of extraction buffer (50mM HEPES, pH 7.6, 120mM KCl, 2mM Mg(OAc)_2_, 2mM CaCl_2_, 2mM Dithiothreitol, 1mg/mL heparin, 0.1mM Benzamidine, 0.1mg/ml soybean trypsin inhibitor and 0.5mM PMSF). The mixture was transferred to RNAse-free Oak Ridge high-speed centrifuge tubes (ThermoFisher Scientific) and centrifuged at 20,000 rpm for 20 min at 4 °C in JA30.50 Ti fixed angle rotor using Avanti JXN-30 ground centrifuge machine. The supernatant was passed through a cheesecloth, and the filtrate was centrifuged again to remove the remaining cell debris. The supernatant was collected and passed through a 0.45 µm syringe filter, and the filtrate was layered on top of a 20% sucrose cushion (6ml) made in extraction buffer and centrifuged at 52,000 rpm for 8 hrs at 4°C in Ti-70 fixed angle rotor to pellet down 80S ribosome particles. The supernatant was discarded, and the 80S pellet was dissolved in a high salt buffer (50mM HEPES/KOH, pH – 7.6, 500mM KCl, 2mM Mg(OAc)_2_, 2mM CaCl_2_).

About 200 µl of 80S ribosome (A_260_ = 200) was layered onto 15-30% sucrose gradient prepared in high salt buffer and centrifuged at 25,000 rpm for 12 hrs at 4°C in SW28 tubes using SW28 Ti swinging bucket rotor, and the fractions were analysed on the agarose gel. The fractions containing the 80S ribosomes were pooled and pelleted by centrifugation at 52,000 rpm for 8 hours at 4°C through 6ml of 20% sucrose cushion in a Ti-70 fixed angle rotor. The pellet was redissolved in dissociation buffer (150mM KCl, 1mM Mg(OAc)_2_, 0.1mM EDTA, 6mM 2-mercaptoethanol, 50mM Tris-HC1, (pH 7.7), containing 5% sucrose) for dissociation of the 80S into 40S and 60S subunits and then treated with 2mM Puromycin for 10 min at 4°C followed by 15 minutes at 37°C. The 500 µl of redissolved 80S was loaded onto a 10%-35% sucrose gradient and centrifuged for 16 hours at 28,000 rpm at 4°C. The fractions were analysed on agarose gel, and the fractions containing only 40S and 60S were pooled separately and pelleted down at 52,000 rpm for 8 hours at 4°C using a Ti-70 fixed-angle rotor. The pellets were resuspended in the storage buffer (50 mM HEPES (pH 7.6), 150 mM KCl, 10 mM Mg(OAc)_2_, 250 mM Sucrose and 2 mM DTT). For reconstitution of 80 ribosomes, the 40S and 60S subunits were mixed in a 1:1 ratio and incubated for 30 minutes in reconstitution buffer (50 mM HEPES (pH 7.6)), 150 mM K(OAc), 20mM Mg(OAc)_2_, 2mM DTT). The reaction was layered onto a 15-30% sucrose density gradient and centrifuged at 25,000 rpm for 12 hours at 4°C in SW-28 tubes using an SW-28 swinging bucket rotor. The fractions were visualized on agarose gel, and the fractions containing 80S were pooled and centrifuged at 52,000 rpm for 8 hours at 4°C using a Ti-70 fixed angle rotor. The final 80S pellet was resuspended in storage buffer-II (50mM HEPES (pH 7.6)), 150mM K(OAc), 20mM Mg(OAc)_2_, 2 mM DTT), and used for Cryo-EM data collection.

### Cryo-EM grid preparation, data collection and processing

3µl of 70nM 80S ribosome was added to a glow-discharged carbon-coated Quantifoil R 2/2 Holey carbon copper grid. After blotting for 3.5 sec and 10 sec wait time in Vitrobot Mark IV at 16 °C and 100% humidity, the sample was plunged into liquid ethane. Cryo-EM data were collected on FEI Titan Krios G3 transmission electron microscope equipped with a FEG at 300 keV with automated data collection software EPU (Thermo FisherScientific). All data were collected using a Falcon III detector at a nominal magnification of 75,000X and a pixel size of 1.07 Å with a total electron dose of 44.60 e^-^/Å^2^ fractionated over 30 frame movies with a dose rate of ∼1.4 e-/Å^2^/frame.

The images in the first dataset displayed good quality of 60S particles and very few 80S particles. The micrographs were processed in Relion 3.1 to obtain a 2.65 Å resolution map of the 60S. The data processing strategy used is summarized in **(Supp Figure 1A)**. Briefly, 8,323 movies were used for motion correction using Relion’s implementation (Zivanov et al., 2019), followed by CTF estimation using CTFFIND4 (Rohou & Grigorieff, 2015). A total of 17,44,676 particles were picked using automated particle picking (Scheres, 2012) and subjected to unsupervised 2D classification to remove the junk particles. A stack of 17,17,534 particles from clean 60S classes obtained in 2D classification was used for reference-based 3D classification into five different classes. One class containing 7,96,258 particles showed clean 60S particles, and it was autorefined to a global resolution of 2.98Å. The resolution was further improved to 2.65Å after particle CTF refinement and Bayesian polishing as per the gold standard FSC of 0.143 (**Supp Figure 1C**). Local resolution analysis was performed using LocRes, and the final map exhibited a range of resolution from 2.5 Å in the core to 4Å towards the periphery (**Supp Figure 1C**).

The other dataset, which largely consisted of 80S particles, was processed in Cryosparc v3.1.1 (Punjani et al, 2020, 2017; Rubinstein & Brubaker, 2015). The overall strategy for data processing is summarized in (**Supp Figure 1B)**. Briefly, the alignment of the image stacks was performed using Patch Motion Correction, and CTF estimation was performed using Patch CTF estimation. Initially, blob picking was performed to select the particles, which was further used for 2D Classification. The classes obtained were used for template-based particle picking followed by 2D Classification to remove junk particles. Clean 2D Classes were subjected to Ab-initio reconstruction into 5 different classes. Clean 80S classes were used for homogenous refinement, resulting in an overall resolution of 2.71 Å for the complete 80S map (**Supp Figure 1D**) To improve it further, local refinements using focused masks were performed on the 60S and 40S subunits independently, which increased the resolution to 2.69 Å (**Supp Figure 1E**) and 2.88 Å, (**Supp Figure 1F**) respectively. A focus refinement over the 40S body resulted in a resolution of 2.84 Å (**Supp Figure 1G)**. Further, we observed that the quality of density of the 40S head region was better in the whole 40S map than the focused refined maps of the head alone. Therefore, model building for the 40S head region was done in the 40S map.

### Model building and refinement

Initially, the 80S ribosome structure from wheat at low resolution (PDB ID 4V7E) (Gogala *et al*, 2014) was used as a template for model building. Briefly, the atomic coordinates were rigid body fit into the 80S map using the dock in map followed by real-space refinement modules in Phenix (Adams *et al*, 2010; Terwilliger *et al*, 2020; Liebschner *et al*, 2019). Model building was performed by an iterative cycle of manual building in Coot (Emsley & Cowtan, 2004) and real space refinement in Phenix, which significantly improved the model’s geometry and fit in the map. Blurred maps of different B-factors were prepared using MRC to MTZ module of CCPEM (Wood *et al*, 2015) to perform model building into the low-resolution peripheral regions of the ribosome.

Later, when high-resolution structures of tomato ribosome (Cottilli *et al*., 2022) (PDB IDs 7QIW, 7QIX & 7QIY for the 60S, 40S body & 40S head region) became available, we used these structures for the model building of the missing regions and validation and/or identification of post-translational modifications and metal ions such as K^+^ ion and Mg^2+^. Model building was followed by validation using the MolProbity module in Phenix, and the figures were prepared using Chimera (Hertig *et al*, 2015; Pettersen *et al*, 2004), ChimeraX (Pettersen *et al*, 2021) and PyMol (DeLano, W.L. (2002). The PyMOL molecular graphics system on world wide web. - References - Scientific Research Publishing).

### Sequence alignment and Phylogenetic analysis

Sequences used for analysis were obtained from NCBI sequence database. The accession IDs for the sequences are listed in Supplementary Tables (Table 2 – Table 8). Sequence alignments were performed in Clustal Omega (Sievers *et al*, 2011). For phylogenetic analysis, sequences were aligned using MUSCLE (Edgar & Batzoglou, 2006) and Jalview (Waterhouse *et al*, 2009).

## Acknowledgements

R.K.M. and P.S. acknowledge the Indian Institute of Science for Ph.D. fellowships. We thank the InSTEM National Cryo-EM facility for data collection. We are grateful to Sucharita Bose for providing technical help with Cryo-EM. We thank the Beagle Computation Cluster team for managing the computational platform, which was used for data processing of the 80S dataset. The authors acknowledge laboratory colleagues for their critical comments on the manuscript.

## Author’s Contribution

R.K.M., and P.S. purified ribosomes and prepared Cryo-EM samples, R.K.M. and A.B.U. processed the data, R.K.M., P.S. and F.T.K. performed model building, R.K.M., and P.S. analysed the data and prepared the figures; T.H. supervised the work and helped to write the manuscript.

## Data availability statement

Cryo-EM maps of wheat 80S, 60S, 40S and 40S body have been deposited in the EMDB with accession codes EMD-XXXX, EMD-XXXX, EMD-XXXX and EMD-XXXX, respectively.

Atomic coordinates of the refined 60S and 40S have been deposited in PDB with accession codes XXXX and XXXX, respectively.

