## Supplementary Figures for "Atomic structure of wheat ribosome reveals unique features of the plant ribosomes"

**This file contains Supplementary Information: 8 Supp Figures and 8 Tables**

### Supp Fig 1

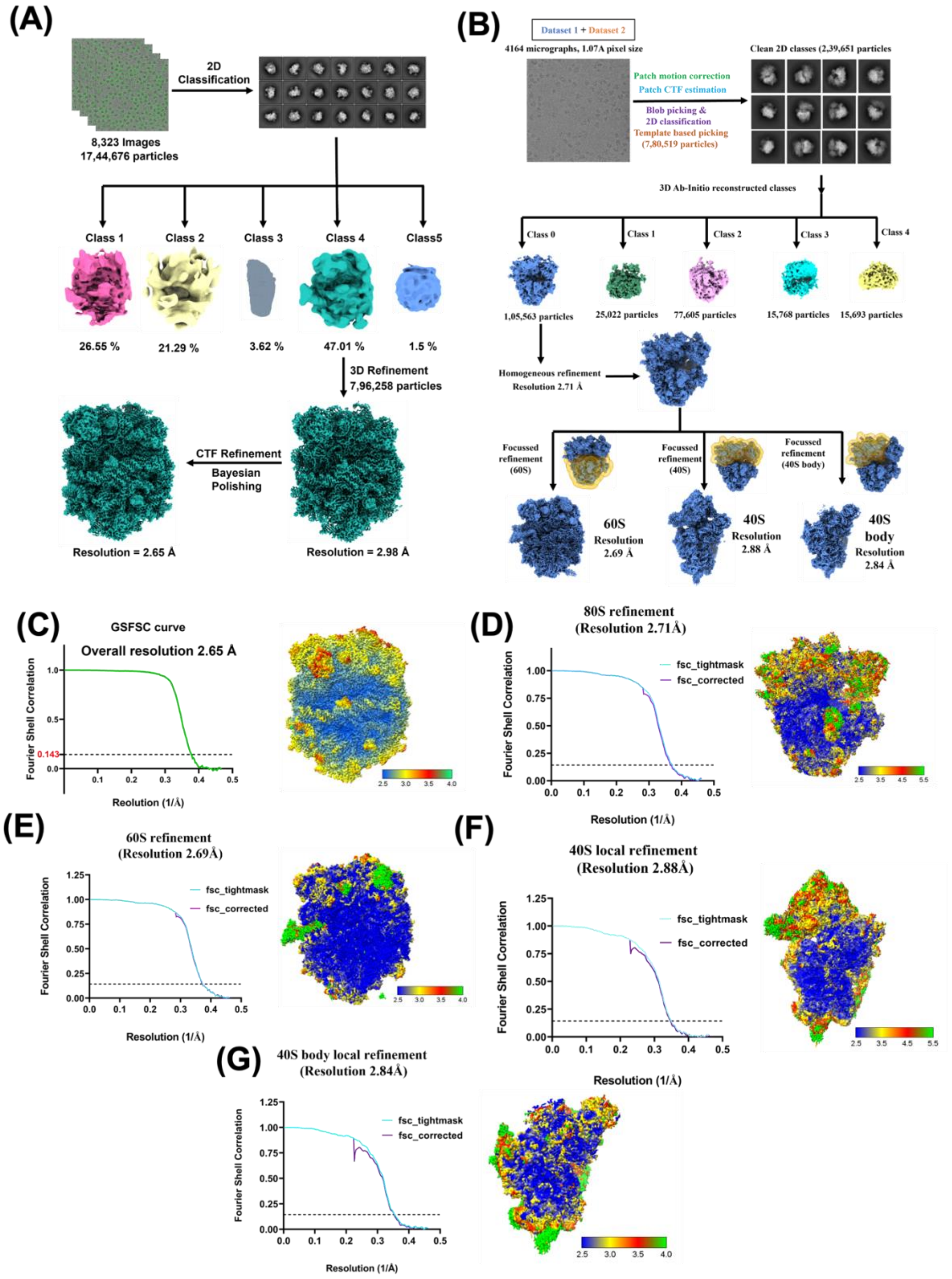


**Supp Figure 1** Cryo-EM data processing for the wheat ribosome

**(A)** Data processing pipeline followed for the 60S dataset. The dataset of 8,323 movies was used and 17,44,676 particles were selected and subjected to 2D classification and reference-based 3D classification leading to the removal of more than 50% of the junk particles. Class 4 obtained in 3D classification was used for 3D refinement resulting in a resolution of 2.98Å after postprocessing, followed by per-particle CTF refinement and Bayesian polishing. The polished particles were used for further 3D refinement leading to a global resolution of 2.65 Å after postprocess.

**(B)** Data processing strategy followed for the 80S dataset: A total of 4,164 movies were collected and were subjected to patch motion correction and CTF estimation, and a total of 7,80,519 particles were picked using blob picking and subjected to multiple rounds of 2D classification to remove the junk particles; then an Ab-initio reconstruction into 5 classes was performed, and the class containing clean 80S was used for the final homogeneous refinement resulting into a resolution of 2.71Å followed by focused refinement on the 60S, 40S subunit and 40S body resulting into a resolution of 2.69Å, 2.88Å and 2.84Å respectively as depicted by the FSC curve and local resolution maps for Cryosparc data processing

**(C)** Resolution estimation using Fourier Shell Correlation (FSC) plot for the autorefined map of the 60S dataset and Local resolution map for the 60S ribosome from wheat showing a range of resolution from 2.5Å in the core to the surface of the ribosome to 3.5Å in the peripheral region

**(D)** Resolution estimation using FSC curve for the homogenous refined map of the 80S dataset and Local resolution map for the 80S ribosome from wheat showing a range of resolution from 2.5Å in the core to 5.5Å or more in the peripheral region

**(E)** Resolution estimation using FSC curve for the local refined map of the 60S and Local resolution map for the 60S ribosome from wheat showing a range of resolution from 2.5Å in the core to 3.5Å or more in the peripheral region

**(F)** Resolution estimation using FSC curve for the local refined map of the 40S and Local resolution map for the 40S ribosome from wheat showing a range of resolution from 2.5Å in the core to 3.5Å or more in the peripheral region

(G) Resolution estimation using FSC curve for the local refined map of the 40S body and Local resolution map for the 40S body showing a range of resolution from 2.5Å in the core to 3.5Å or more in the peripheral region

### Supp Fig 2

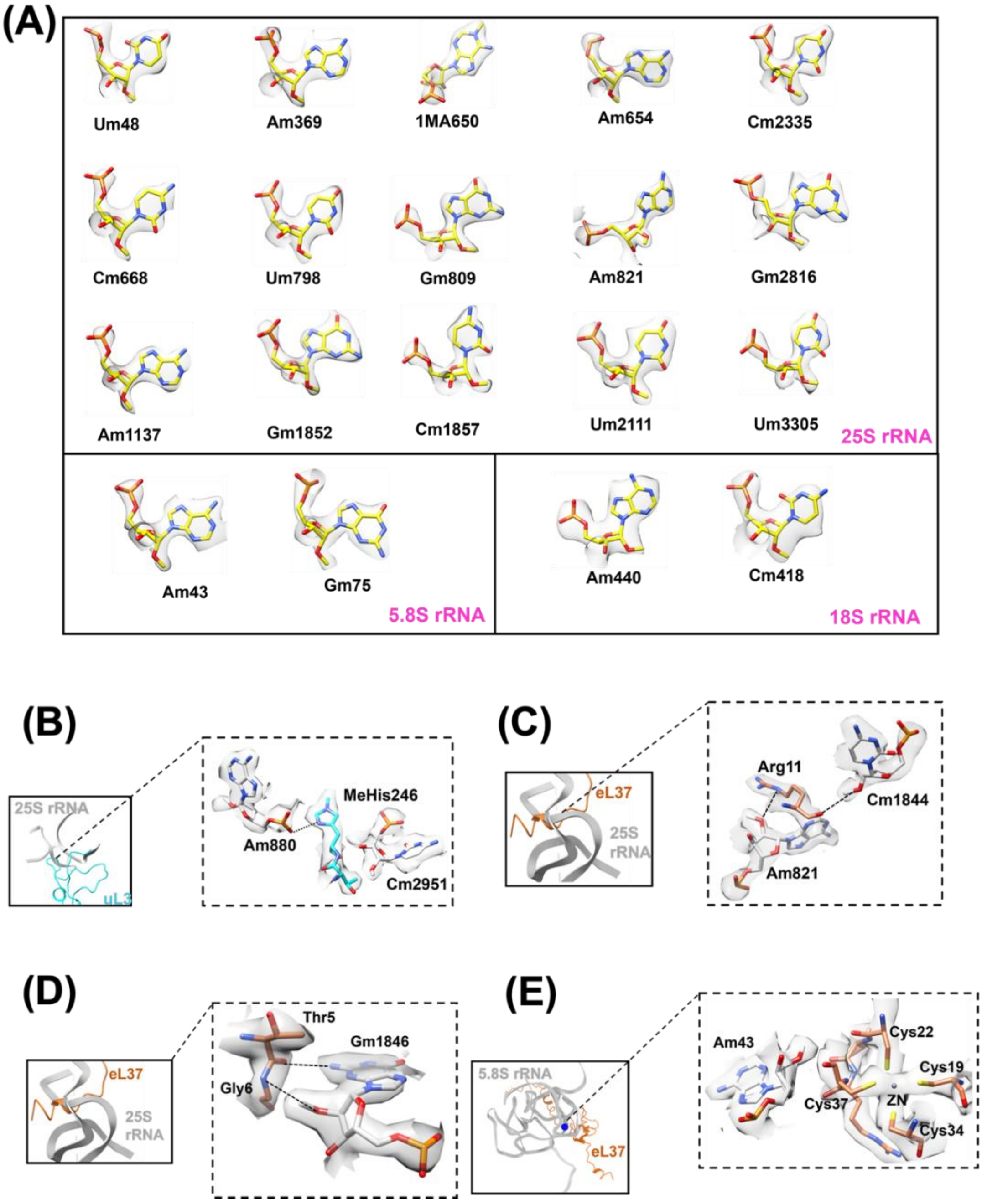


**Supp Figures 2** The density for the methylation in the cryo-EM map of wheat ribosome and common features between wheat and tomato ribosome

(A) Density render diagrams for representative nucleotides that reflect the fit of chemical modification in the map
(B) Interaction of universally conserved modified amino acid His246 with 25S rRNA bases harbouring plant specific 2’O-methylation (Am880 and Cm2951)
(C) Plant specific Gm1846 forming interactions with N-terminal region of eL37
(D) Arg11 of eL37 RP forms interactions with bases Cm1844 and Am821, which possess 2’O-methylation only in plants
(E) Am43 in plant 5.8S rRNA directly interact with Zinc-finger motif of eL37

### Supp Fig 3

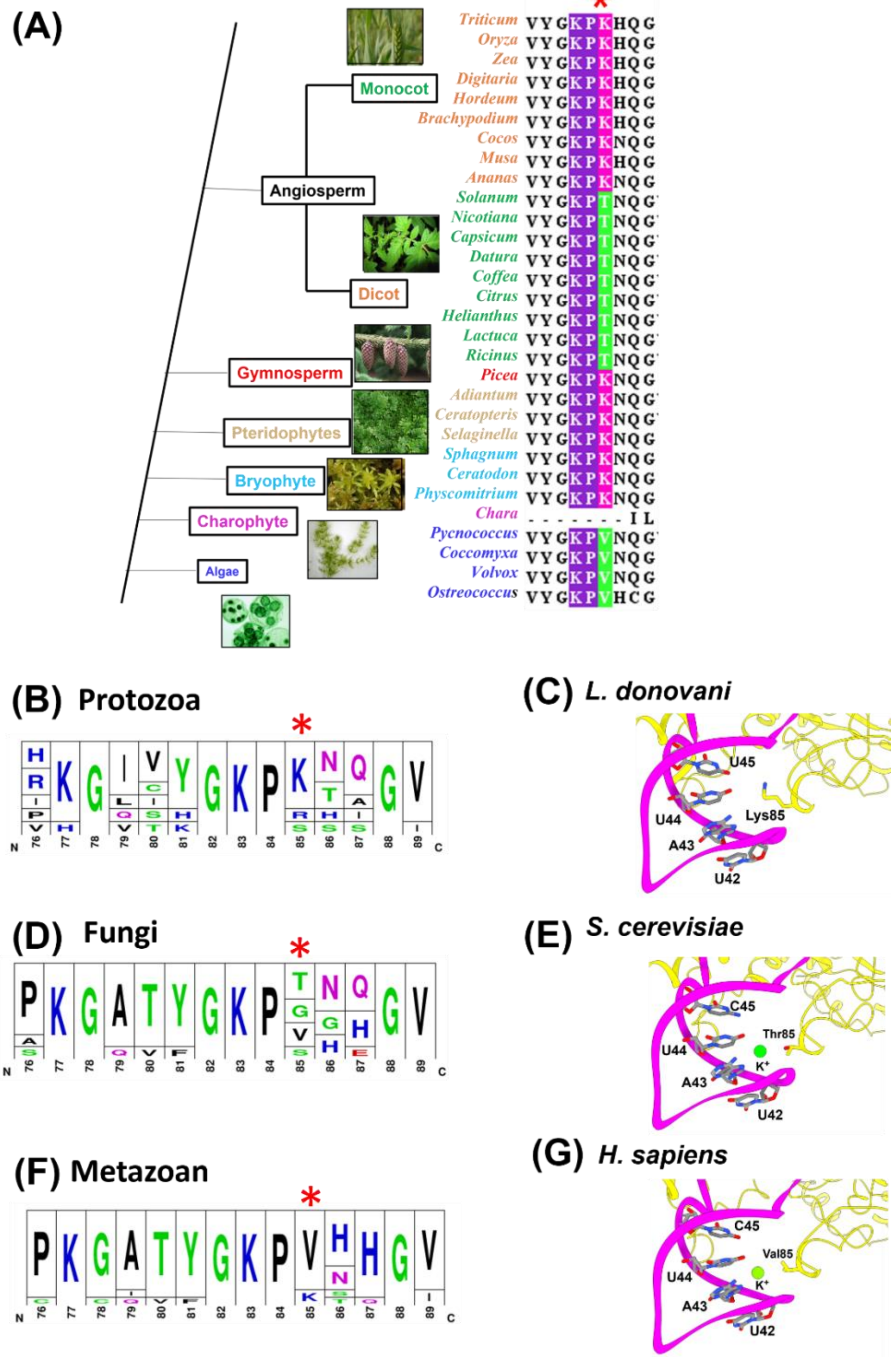


**Supp Figures 3** Sequence and structure analysis of eL15 from different species

(A) The sequence analysis in plants shows the Lys85 (shown by an asterisk) to be highly conserved in Bryophyte, Pteridophyte, Gymnosperm and Angiosperm

(B) & (C) Sequence analysis of eL15 and structural analysis of eL15-H11 interface from protozoa

(D) & (E) Sequence analysis of eL15 and structural analysis of eL15-H11 interface from fungi

(F) & (G) Sequence analysis of eL15 and structural analysis of eL15-H11 interface from metazoan

### Supp Fig 4

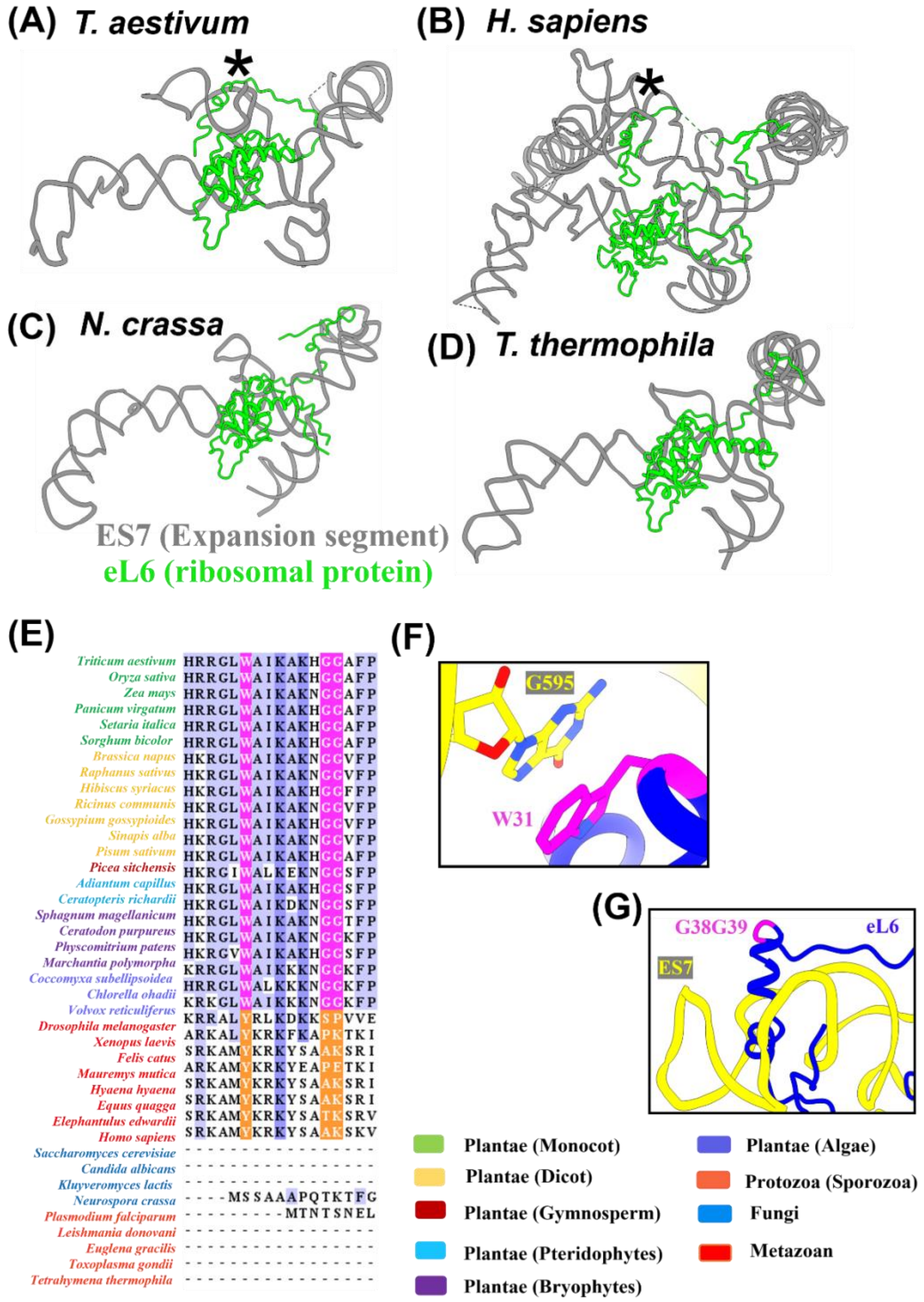


**Supp Figure 4** Interaction between eL6 and Expansion segment 7$^{c}$ (ES7$^{c}$)

(A) & (B) Intercalation of NTT of eL6 through ES7$^{c}$ in (A) plant (wheat) and (B) metazoan (human), respectively, as depicted by an asterisk to represent the position of intercalation

(C) & (D) No intercalation through ES7$^{c}$ by eL6 observed in (C) fungi and (D) protozoan, respectively

(E) Sequence comparison of NTT of plant eL6 with protozoa, yeast, and metazoan

(F) Stacking interaction between a plant-specific W31 in eL6 with G595 of 25S rRNA

(G) Plant-specific Glycine dipeptide G38G39 depicted in magenta, facilitates the bending of the eL6-NTT towards ES7$^{c}$

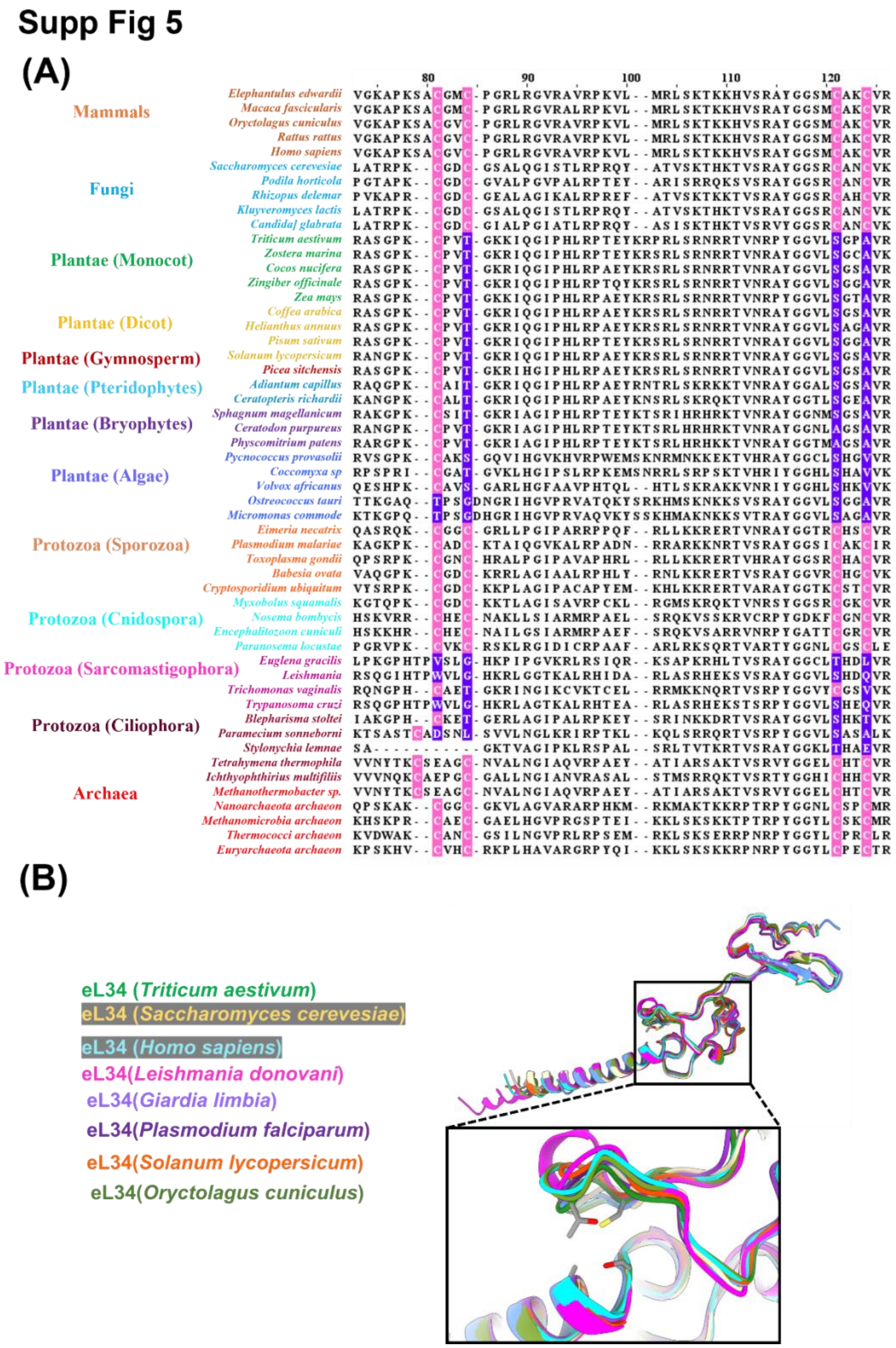


**Supp Figure 5** Sequence alignment of eL34 from various eukaryotes

(A) Sequence comparison shows an absence of the otherwise conserved cysteine in the case of plants and protozoa

(B) Structure comparison of eL34 from species with and without Zinc-finger motif showing no overall difference in the conformation

### Supp Fig 6

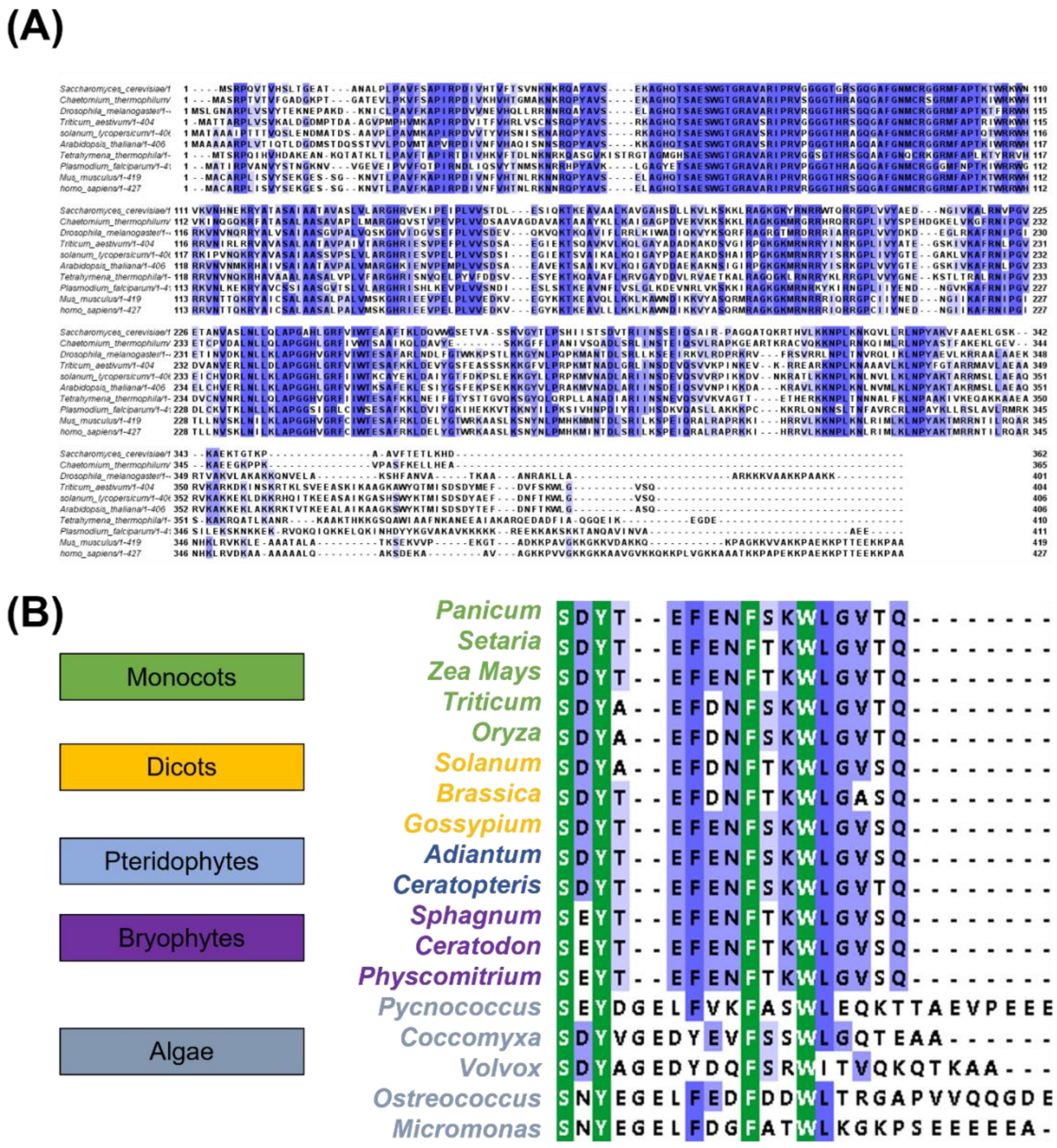


**Supp Figure 6** Sequence analysis of uL4 from plants and other species

(A) Multiple sequence alignment of uL4 from different eukaryotic organisms showing varying lengths of uL4 CTT

(B) Multiple sequence alignments of uL4 CTT from higher plants to lower algae showing conservation of residues (highlighted in green) involved in the interaction with neighbouring eL20 and eL21

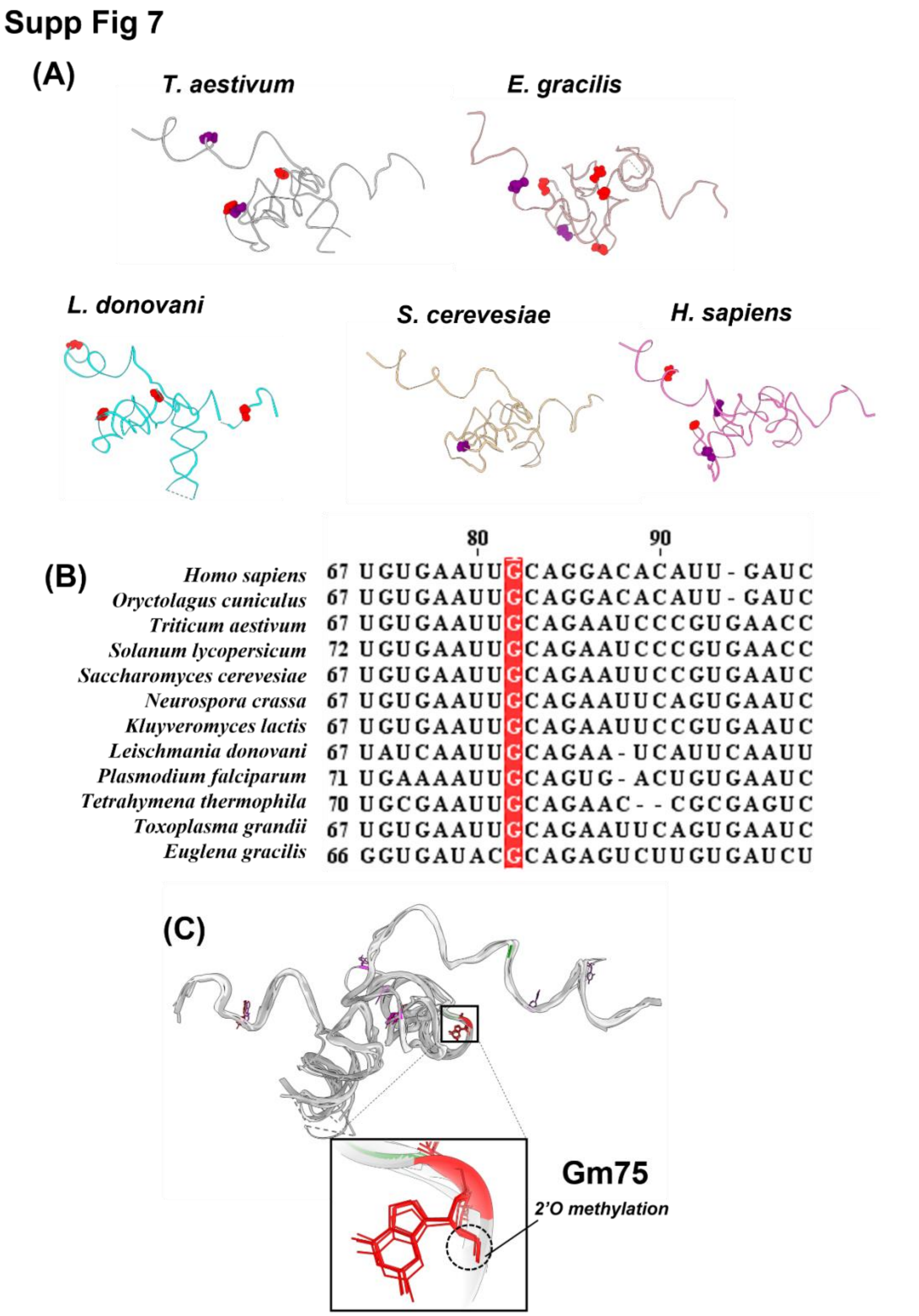


**Supp Figure 7** Potential role of conserved Gm75 in translation

(A) Chemical modifications on 5.8S rRNA of different species is represented where the position as well as number of modification varies across eukaryotes

(B) Sequence comparison of 5.8S rRNA from different species shows high conservation of Guanine at this position throughout evolution

(C) Superposition of the 5.8S rRNA shows the conservation of modification on G75 in different species (yeast ribosome shows no modification (PDB-ID: 4V88)

### Supp Fig 8

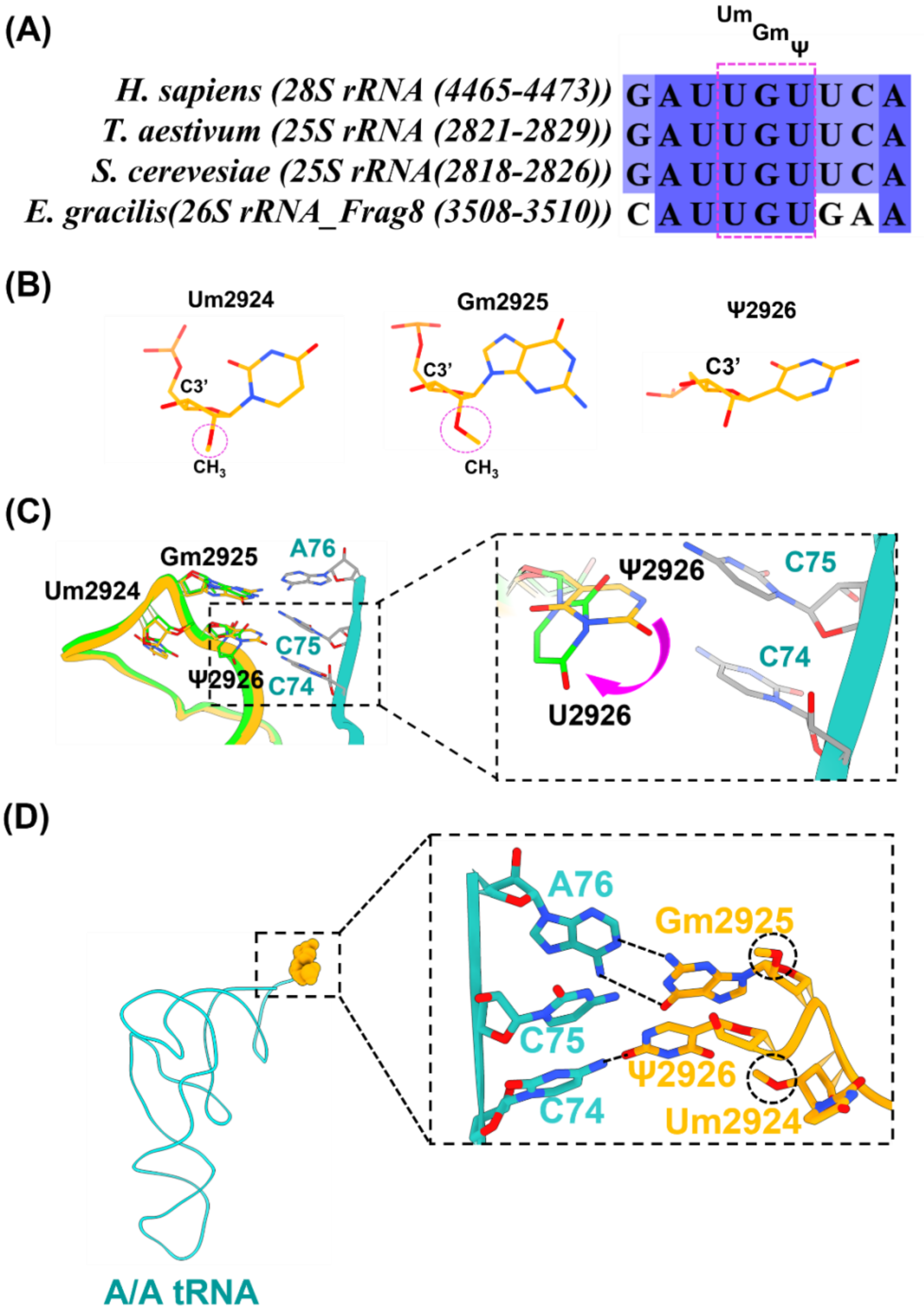


**Supp Figure 8** Conservation of modified triplet and potential of 2'O ribose methylation to induce C3' endo conformation

(A) Sequence comparison shows the conservation of modified nucleotides UmGmΨ triplet (Um2924Gm2925Ψ2926) through eukaryotic evolution

(B) A C3' endo conformation is induced in Um2924Gm2925Ψ2926 of H92 by the presence of methylation at 2'O ribose of the UmGmΨ triplet

(C) Comparison of H92 between wheat ribosome structure (orange) and pseudouridine-less ribosome structure (green) from yeast (PDB ID 7MPI) showing the altered conformation of U2926

(D) Planar arrangement of the modified residues Um2924Gm2925Ψ2926 involved in interaction with CCA at the acceptor arm of tRNA

**Table 1: Data Collection and refinement statistics**

| **Data collection and processing** | **60S subunit** | **40S subunit** |
|---|---|---|
| Microscope | Titan Krios | Titan Krios |
| Voltage | 300 kV | 300 kV |
| Magnification | 75,000x | 75,000x |
| Detector | Falcon III | Falcon III |
| Sampling (Å/pixel) | 1.07 | 1.07 |
| Total electron dose [$e^-$/Å$^2$] | 44.60 | 44.60 |
| Symmetry imposed | C1 | C1 |
| Average resolution (Å) | 2.7 | 2.9 |
| **Atomic Model composition** | | |
| Chains RNA/Protein | 3/44 | 1/31 |
| Non-hydrogen atoms | 116711 | 69046 |
| Amino Acids | 6191 | 4453 |
| Nucleotides | 3134 | 1571 |
| Number of ligand atoms | 317 | 87 |
| $Zn^{2+}$/$Mg^{2+}$/$K^+$ | 3/242/72 | 1/70/16 |
| **Refinement Statistics** | | |
| Model to map CC ($CC_{mask}$/$CC_{box}$/$CC_{peaks}$/$CC_{volume}$) | 0.82/0.73/0.72/0.82 | 0.77/0.67/0.76/0.72 |
| Resolution [Å] by model-to-map FSC, threshold 0.50 (masked/unmasked) | 2.6/2.5 | 3.3/3.4 |
| Bond length rmsd | 0.006 | 0.003 |
| Bond angle rmsd | 0.696 | 0.636 |
| **Validation** | | |
| Clash score | 6.57 | 12.43 |
| Rotamer outliers [%] | 3.32 | 0.47 |
| Ramachandran plot [%] (Favored/allowed/disallowed) | 95.76/4.24/0 | 93.56/6.35/0.09 |
| CaBLAM outliers [%] | 2.32 | 3.50 |
| MolProbity Score | 2.32 | 2.03 |
| **Accession ID** | | |
| EMDB ID | | |
| PDB ID | | |

**Table 2:Accession IDs for eL15 ribosomal protein**

| Organism | Accession ID |
|---|---|
| *Triticum aestivum* | XP_044323743.1 |
| *Oryza brachyantha* | XP_040380504.1 |
| *Zea mays* | NP_001150386.2 |
| *Solanum lycopersicum* | XP_004241917.1 |
| *Nicotiana tomentosiformis* | XP_009601449.1 |
| *Capsicum annuum* | XP_016582172.1 |
| *Picea sitchensis* | ABK22257.1 |
| *Adiantum capillus* | KAI5058505.1 |
| *Ceratopteris richardii* | KAH7288671.1 |
| *Selaginella moellendorffii* | XP_002961462.1 |
| *Sphagnum magellanicum* | KAH9540499.1 |
| *Ceratodon purpureus* | KAG0554928.1 |
| *Physcomitrium patens* | XP_024357573.1 |
| *Coccomyxa subellipsoidea* | XP_005646118.1 |
| *Volvox africanus* | GIL45611.1 |
| *Cyanidioschyzon merolae* | XP_005534852.1 |
| *Plasmodium vivax* | XP_001615040.1 |
| *Leishmania donovani* | XP_003863082.1 |
| *Euglena gracilis* | QLA09621.1 |
| *Giardia intestinalis* | XP_001708727.1 |
| *Paramecium octaurelia* | CAD8175268.1 |
| *Toxoplasma gondii* | XP_002366469.1 |
| *Stentor coeruleus* | OMJ69253.1 |
| *Saccharomyces cerevisiae* | AJS80338.1 |
| *Kluyveromyces lactis* | QEU62382.1 |
| *Candida viswanathii* | RCK59573.1 |
| *Rhizopus arrhizus* | KAG0748686.1 |
| *Podila verticillata* | KAF9367241.1 |
| *Rhizoctonia solani* | CAE6457639.1 |
| *Cladochytrium replicatum* | KAI8800011.1 |
| *Lobosporangium transversale* | XP_021882443.1 |
| *Homo sapiens* | NP_001240308.1 |
| *Oryctolagus cuniculus* | XP_002716262.1 |
| *Bos taurus* | DAA16199.1 |
| *Xenopus laevis* | XP_018124768.1 |
| *Chelydra serpentina* | KAG6940002.1 |
| *Mauremys mutica* | XP_044863280.1 |
| *Gallus gallus* | NP_001292094.1 |
| *Anas platyrhynchos* | XP_005029295.3 |
| *Hydra vulgaris* | P61368.2 |
| *Periplaneta americana* | KAJ4450415.1 |
| *Caenorhabditis elegans* | NP_499964.1 |

**Table 3: Accession IDs for eL6 ribosomal protein sequences**

| Organism | Accession ID |
|---|---|
| *Triticum aestivum* | XP_044403216.1 |
| *Oryza sativa* | XP_015627578.1 |
| *Zea mays* | NP_001130513.1 |
| *Panicum virgatum* | XP_039847586.1 |
| *Setaria italica* | XP_004975938.1 |
| *Sorghum bicolor* | XP_002448023.1 |
| *Brassica napus* | XP_013664611.2 |
| *Raphanus sativus* | XP_018433501.1 |
| *Hibiscus syriacus* | XP_039007682.1 |
| *Ricinus communis* | XP_002513192.1 |
| *Gossypium gossypioides* | MBA0750743.1 |
| *Sinapis alba* | KAF8114319.1 |
| *Pisum sativum* | KAI5405181.1 |
| *Picea sitchensis* | ABK21588.1 |
| *Adiantum capillus* | KAI5075325.1 |
| *Ceratopteris richardii* | KAH7291547.1 |
| *Sphagnum magellanicum* | KAH9535645.1 |
| *Ceratodon purpureus* | KAG0570454.1 |
| *Physcomitrium patens* | XP_024369198.1 |
| *Marchantia polymorpha* | OAE31965.1 |
| *Coccomyxa subellipsoidea* | XP_005647799.1 |
| *Chlorella ohadii* | KAI7837103.1 |
| *Volvox reticuliferus* | GIL85813.1 |
| *Felis catus* | XP_006938548.1 |
| *Hyaena hyaena* | XP_039102100.1 |
| *Equus quagga* | XP_046497158.1 |
| *Elephantulus edwardii* | XP_006901041.1 |
| *Homo sapiens* | 6QZP (PDB) |
| *Mauremys mutica* | XP_044845807.1 |
| *Xenopus laevis* | XP_018119120.1 |
| *Drosophila melanogaster* | NP_651876.1 |
| *Candida albicans* | AJV67506.1 |
| *Kluyveromyces lactis* | XP_451742.1 |
| *Neurospora crassa* | CAE76504.1 |
| *Plasmodium falciparum* | XP_001350109.1 |
| *Leishmania donovani* | 3JCS (PDB ID) |
| *Euglena gracilis* | QLA09617.1 |
| *Toxoplama gondii* | PIM00564.1 |
| *Tetrahymena thermophila* | XP_001019693.2 |

**Table 4: Accession IDs for uL22 ribosomal Protein**

| Organism | Accession ID |
|---|---|
| *Triticum aestivam* | XP_044431146.1 |
| *Oryza sativa* | XP_015612256.1 |
| *Solanum lycopersicum* | XP_004236338.1 |
| *Arabidopsis thaliana* | NP_174060.1 |
| *Saccharomyces cerevisiae* | AJS47412.1 |
| *Candida albicans* | XP_712916.1 |
| *Drosophila melanogaster* | NP_001284961.1 |
| *Caenorhabditis elegans* | NP_740781.1 |
| *Homo sapiens* | KAI2586824.1 |
| *Elephantulus edwardii* | XP_006903542.1 |
| *Plasmodium falciparum* | XP_001350227.1 |
| *Euglena gracilis* | QLA09612.1 |
| *Escherichia coli* | WP_266146827.1 |
| *Salmonella enterica* | NRK38637.1 |

**Table 5: Interacting residues at eL34 and 28S rRNA interface**

| eL34 residues | 25S rRNA residues |
|---|---|
| K49 | C1708 |
| K50 | A1706 |
| Q52 | A1707 |
| G53 | G1639 (through $K^+$)* |
| R58 | G1591 |
| H56 | A1740 |
| R56 | A1655 |
| T60 | A1592 |
| K63 | G1616 |
| R64 | U1822 |
| R66 | A1593 |
| S68 | A1642 |
| R69 | G1644 |
| N70 | G1639 (through $K^+$)* |
| R71 | C1739 |
| R72 | U1822 |
| N75 | C1632 |
| R76 | C1638 |

*The interaction between the eL34 residues and rRNA at these positions is coordinated through $K^+$ ion

**Table 6: Accession IDs for eL34 ribosomal protein**

| Organism | Accession ID |
|---|---|
| *Elephantulus edwardii* | XP_006884412 |
| *Macaca fascicularis* | XP_045253972.1 |
| *Oryctolagus cuniculus* | PDB: 7NFX_g |
| *Rattus rattus* | CAA32574.1 |
| *Homo sapiens* | NP_000986.2 |
| *Saccharomyces cerevesiae* | PDB: 4V6I_Bi |
| *Podila horticola* | KAG0023736.1 |
| *Rhizopus delemar* | EIE75655.1 |
| *Kluyveromyces lactis* | NP_010977.2 |
| *Candida glabrata* | XP_449882.1 |
| *Triticum aestivum* | AAW50987.1 |
| *Cocos nucifera* | XP_008802610.1 |
| *Zingiber officinale* | XP_042388246.1 |
| *Zea mays* | NP_001132454.1 |
| *Coffea arabica* | XP_027075604.1 |
| *Helianthus annuus* | KCW64962.1 |
| *Pisum sativum* | KAI5420024.1 |
| *Solanum lycopersicum* | XP_004238799.1 |
| *Picea sitchensis* | ABK21342.1 |
| *Adiantum capillus* | KAI5062804.1 |
| *Ceratopteris richardii* | KAH7284748.1 |
| *Sphagnum magellanicum* | KAH9559090.1 |
| *Ceratodon purpureus* | KAG0605459.1 |
| *Physcomitrium patens* | XP_024360062.1 |
| *Pycnococcus provasolii* | GHP03976.1 |
| *Coccomyxa sp* | BDA42094.1 |
| *Volvox africanus* | GIL50355.1 |
| *Ostreococcus tauri* | XP_022838836.1 |
| *Micromonas commoda* | XP_002504814.1 |
| *Euglena gracilis* | QLA09635.1 |
| *Leishmania donovani* | XP_001686957.1 |
| *Trichomonas vaginalis* | XP_001303734.1 |
| *Trypanosoma cruzi* | XP_815813.1 |
| *Eimeria necatrix* | XP_013335039.1 |
| *Plasmodium malariae* | SBT74739.1 |
| *Toxoplasma gondii* | XP_002366393.1 |
| *Babesia ovata* | XP_028868707.1 |
| *Cryptosporidium ubiquitum* | XP_028875035.1 |
| *Myxobolus squamalis* | KAF1744176.1 |
| *Nosema bombycis* | ADZ95696.1 |
| *Encephalitozoon cuniculi* | NP_597582.1 |

| | |
|---|---|
| *Paranosema locustae* | PDB: 6ZU5_LGG |
| *Tetrahymena thermophila* | XP_001021598.1 |
| *Blepharisma stoltei* | CAG9333327.1 |
| *Paramecium sonneborni* | CAD8077493.1 |
| *Ichthyophthirius multifilii* | XP_004037728.1 |
| *Stylonychia lemnae* | CDW83538.1 |
| *Methanothermobacter sp.* | XP_001021598.1 |
| *Nanoarchaeota sp.* | MCK4589265.1 |
| *Methanomicrobia sp.* | MCD6128027.1 |
| *Thermococci sp.* | RLF91603.1 |
| *Euryarchaeota sp.* | MBM4240109.1 |

**Table 7: Accession IDs for uL4 ribosomal protein**

| Organism | Accession ID |
|---|---|
| *Panicum miliaceum* | RLM84596.1 |
| *Setaria italica* | XP_004981434.1 |
| *Zea mays* | ACG25262.1 |
| *Oryza sativa Japonica Group* | XP_015647011.1 |
| *Triticum aestivum* | 4V7E_CC |
| *Brassica rapa* | XP_009146912.2 |
| *Gossypium arboreum* | XP_017649246.1 |
| *Adiantum capillus-veneris* | KAI5082907.1 |
| *Ceratopteris richardii* | KAH7365535.1 |
| *Sphagnum magellanicum* | KAH9563930.1 |
| *Ceratodon purpureus* | KAG0597514.1 |
| *Physcomitrium patens* | XP_024362557.1 |
| *Pycnococcus provasolii* | GHP11437.1 |
| *Ostreococcus tauri* | XP_022840336.1 |
| *Volvox africanus* | GIL42455.1 |
| *Coccomyxa sp. Obi* | BDA44074.1 |
| *Micromonas commoda* | XP_002506564.1 |
| *Homo sapiens* | NP_000959.2 |
| *Mus musculus* | NP_077174.1 |
| *Drosophila melanogaster* | NP_524538.2 |
| *Arabidopsis thaliana* | NP_187574.1 |
| *Tetrahymena thermophila* | XP_001017488.2 |
| *Plasmodium falciparum* | XP_001351629.1 |
| *Saccharomyces cerevisiae* | NP_009587.1 |
| *Thermochaetoides thermophila* | XP_006696470.1 |

**Table 8: Source of the sequences of 5.8S rRNA**

| Organism | Source of Sequence |
|---|---|
| *Triticum aestivum* | 4V7E |
| *Solanum lycopersicum* | 7QIW |
| *Homo sapiens* | 6QZP |
| *Oryctolagus cuniculus* | 7O7Y |
| *Saccharomyces cerevesiae* | 4V88 |
| *Neurospora crassa* | 6YWS |
| *Kluyveromyces lactis* | 6UZ7 |
| *Plasmodium falciparum* | 5UMD |
| *Tetrahymena thermophila* | 4V8P |
| *Toxoplasma gondii* | 5XXB |
| *Euglena gracillis* | 6ZJ3 |
